# Genetic discovery and translational decision support from exome sequencing of 20,791 type 2 diabetes cases and 24,440 controls from five ancestries

**DOI:** 10.1101/371450

**Authors:** Jason Flannick, Josep M Mercader, Christian Fuchsberger, Miriam S Udler, Anubha Mahajan, Jennifer Wessel, Tanya M Teslovich, Lizz Caulkins, Ryan Koesterer, Thomas W Blackwell, Eric Boerwinkle, Jennifer A Brody, Ling Chen, Siying Chen, Cecilia Contreras-Cubas, Emilio Córdova, Adolfo Correa, Maria Cortes, Ralph A DeFronzo, Lawrence Dolan, Kimberly L Drews, Amanda Elliott, James S Floyd, Stacey Gabriel, Maria Eugenia Garay-Sevilla, Humberto García-Ortiz, Myron Gross, Sohee Han, Sarah Hanks, Nancy L Heard-Costa, Anne U Jackson, Marit E Jørgensen, Hyun Min Kang, Megan Kelsey, Bong-Jo Kim, Heikki A Koistinen, Johanna Kuusisto, Joseph B Leader, Allan Linneberg, Ching-Ti Liu, Jianjun Liu, Valeriya Lyssenko, Alisa K Manning, Anthony Marcketta, Juan Manuel Malacara-Hernandez, Angélica Martínez-Hernández, Karen Matsuo, Elizabeth Mayer-Davis, Elvia Mendoza-Caamal, Karen L Mohlke, Alanna C Morrison, Anne Ndungu, Maggie CY Ng, Colm O’Dushlaine, Anthony J Payne, Catherine Pihoker, Broad Genomics Platform, Wendy S Post, Michael Preuss, Bruce M Psaty, Ramachandran S Vasan, N William Rayner, Alexander P Reiner, Cristina Revilla-Monsalve, Neil R Robertson, Nicola Santoro, Claudia Schurmann, Wing Yee So, Heather M Stringham, Tim M Strom, Claudia HT Tam, Farook Thameem, Brian Tomlinson, Jason M Torres, Russell P Tracy, Rob M van Dam, Marijana Vujkovic, Shuai Wang, Ryan P Welch, Daniel R Witte, Tien-Yin Wong, Gil Atzmon, Nir Barzilai, John Blangero, Lori L Bonnycastle, Donald W Bowden, John C Chambers, Edmund Chan, Ching-Yu Cheng, Yoon Cho Shin, Francis S Collins, Paul S de Vries, Ravindranath Duggirala, Benjamin Glaser, Clicerio Gonzalez, Ma Elena Gonzalez, Leif Groop, Jaspal Singh Kooner, Soo Heon Kwak, Markku Laakso, Donna M Lehman, Peter Nilsson, Timothy D Spector, E Shyong Tai, Tiinamaija Tuomi, Jaakko Tuomilehto, James G Wilson, Carlos A Aguilar-Salinas, Erwin Bottinger, Brian Burke, David J Carey, Juliana Chan, Josée Dupuis, Philippe Frossard, Susan R Heckbert, Mi Yeong Hwang, Young Jin Kim, H Lester Kirchner, Jong-Young Lee, Juyoung Lee, Ruth Loos, Ronald CW Ma, Andrew D Morris, Christopher J O’Donnell, Colin NA Palmer, James Pankow, Kyong Soo Park, Asif Rasheed, Danish Saleheen, Xueling Sim, Kerrin S Small, Yik Ying Teo, Christopher Haiman, Craig L Hanis, Brian E Henderson, Lorena Orozco, Teresa Tusié-Luna, Frederick E Dewey, Aris Baras, Christian Gieger, Thomas Meitinger, Konstantin Strauch, Leslie Lange, Niels Grarup, Torben Hansen, Oluf Pedersen, Phil Zeitler, Dana Dabelea, Goncalo Abecasis, Graeme I Bell, Nancy J Cox, Mark Seielstad, Rob Sladek, James B Meigs, Steve Rich, Jerome I Rotter, DiscovEHR Collaboration, CHARGE, LuCamp, ProDiGY, GoT2D, ESP, SIGMA-T2D, T2D-GENES, AMP-T2D-GENES, David Altshuler, Noёl P Burtt, Laura J Scott, Andrew P Morris, Jose C Florez, Mark I McCarthy, Michael Boehnke

## Abstract

Protein-coding genetic variants that strongly affect disease risk can provide important clues into disease pathogenesis. Here we report an exome sequence analysis of 20,791 type 2 diabetes (T2D) cases and 24,440 controls from five ancestries. We identify rare (minor allele frequency<0.5%) variant gene-level associations in (a) three genes at exome-wide significance, including a T2D-protective series of >30 *SLC30A8* alleles, and (b) within 12 gene sets, including those corresponding to T2D drug targets (*p*=6.1×10^−3^) and candidate genes from knockout mice (*p*=5.2×10^−3^). Within our study, the strongest T2D rare variant gene-level signals explain at most 25% of the heritability of the strongest common single-variant signals, and the rare variant gene-level effect sizes we observe in established T2D drug targets will require 110K-180K sequenced cases to exceed exome-wide significance. To help prioritize genes using associations from current smaller sample sizes, we present a Bayesian framework to recalibrate association *p*-values as posterior probabilities of association, estimating that reaching *p*<0.05 (*p*<0.005) in our study increases the odds of causal T2D association for a nonsynonymous variant by a factor of 1.8 (5.3). To help guide target or gene prioritization efforts, our data are freely available for analysis at www.type2diabetesgenetics.org.

## Introduction

To better understand or treat disease, human genetics offers a powerful approach to identify molecular alterations causally associated with physiological traits^1^. Common-variant array-based genome-wide association studies (GWAS) have discovered thousands of genomic loci associated with hundreds of human traits^2^, and further common variant analyses indicate that most complex trait heritability is attributable to modest-effect regulatory variants^3–5^. However, non-coding GWAS associations are challenging to localize to causal variants or genes^6–10^.

Protein-coding variants with strong effects on protein function or disease can offer molecular “probes” into the pathological relevance of a gene^13–15^ and potentially establish a direct causal^16,17^ link between gene gain or loss of function and disease risk^18,19^ – especially when there is evidence of multiple independent variant associations (an “allelic series”) within a gene^18–20^. Several lines of argument^11,12^ predict that strong-effect variants (allelic odds-ratios [OR]>2) will usually be rare (minor allele frequency [MAF]<0.5%) and, in many cases, difficult to accurately study through current GWAS and imputation strategies^13,14^. Whole genome or exome sequencing, by contrast, allows interrogation of the full spectrum of genetic variation.

Previous exome sequencing studies, however, have identified few exome-wide significant rare variant associations^21–26^ for complex diseases such as type 2 diabetes (T2D)^24,27^. This paucity of findings is due in part to the limited sample sizes of previous studies, the largest of which include <10,000 disease cases and fall short of the sample sizes that analytic^12^ and simulation-based calculations^28–30^ predict are needed to identify rare disease-associated variants under plausible disease models. To expand our ability to use rare coding variants to make genetic discoveries and accelerate clinical translation, we collected and analyzed exome sequence data from 20,791 T2D cases and 24,440 controls of multiple ancestries, representing the largest exome sequence analysis to date for T2D.

### Genetic discovery from single-variant and gene-level analysis

Study participants (**Supplementary Table 1**) were drawn from five ancestries (Hispanic/Latino [effective size (N_eff_)=14,442; 33.8%], European [N_eff_=10,517; 24.6%], African-American [N_eff_=5,959; 13.9%], East-Asian [N_eff_=6,010; 14.1%], South-Asian [N_eff_=5,833; 13.6%]) and yielded equivalent statistical power to detect association as a balanced study of ~42,800 individuals or a population-based study (assuming 8% T2D prevalence) of ~152,000 individuals. Power to detect association was improved compared to the previous largest T2D exome sequencing study^24^ of 6,504 cases and 6,436 controls, increasing (for example) from 5% to 90% for a variant with MAF=0.2% and OR=2.5 (**Supplementary Figure 1**).

Exome sequencing to 40x mean depth, variant calling using best-practice algorithms, and extensive data quality control (**Methods; Supplementary Figures 2-5, Supplementary Table 2**) produced a dataset with 6.33M variants, of which 2.3% are common (MAF>5%), 4.2% low-frequency (0.5%<MAF<5%), and 93.5% rare (MAF<0.5%) (**Supplementary Table 3**). These include 2.26M nonsynonymous variants and 871K indels, more than twice the numbers analyzed in the largest previous T2D exome sequencing study^24^.

We first tested whether any of these variants, regardless of allele frequency, exhibited association with T2D (“single-variant” test; **Methods, Supplementary Figure 6**). Based on a previously demonstrated enrichment of coding variants for disease associations^31^, we used an exome-wide significance threshold of *p*=4.3×10^−7^. Eighteen variants (ten nonsynonymous) in seven loci reached this threshold; 13 of these (eight nonsynonymous) reached the traditional genome-wide significance threshold of *p*<5×10^−8^ (**Figure 1a, Supplementary Table 4**). These 18 associations represent a substantial increase over the one association reported from the previous largest T2D exome sequencing study^24^. However, only two of these 18 have not been previously reported by (much larger) GWAS: a variant in *SFI1* (rs145181683, p.Arg724Trp; **Supplementary Figure 7**) that failed to replicate in an independent cohort (N=4,522, p=0.90, **Methods**), and a variant in *MC4R* (rs79783591, p.Ile269Asn).

**Figure 1:**
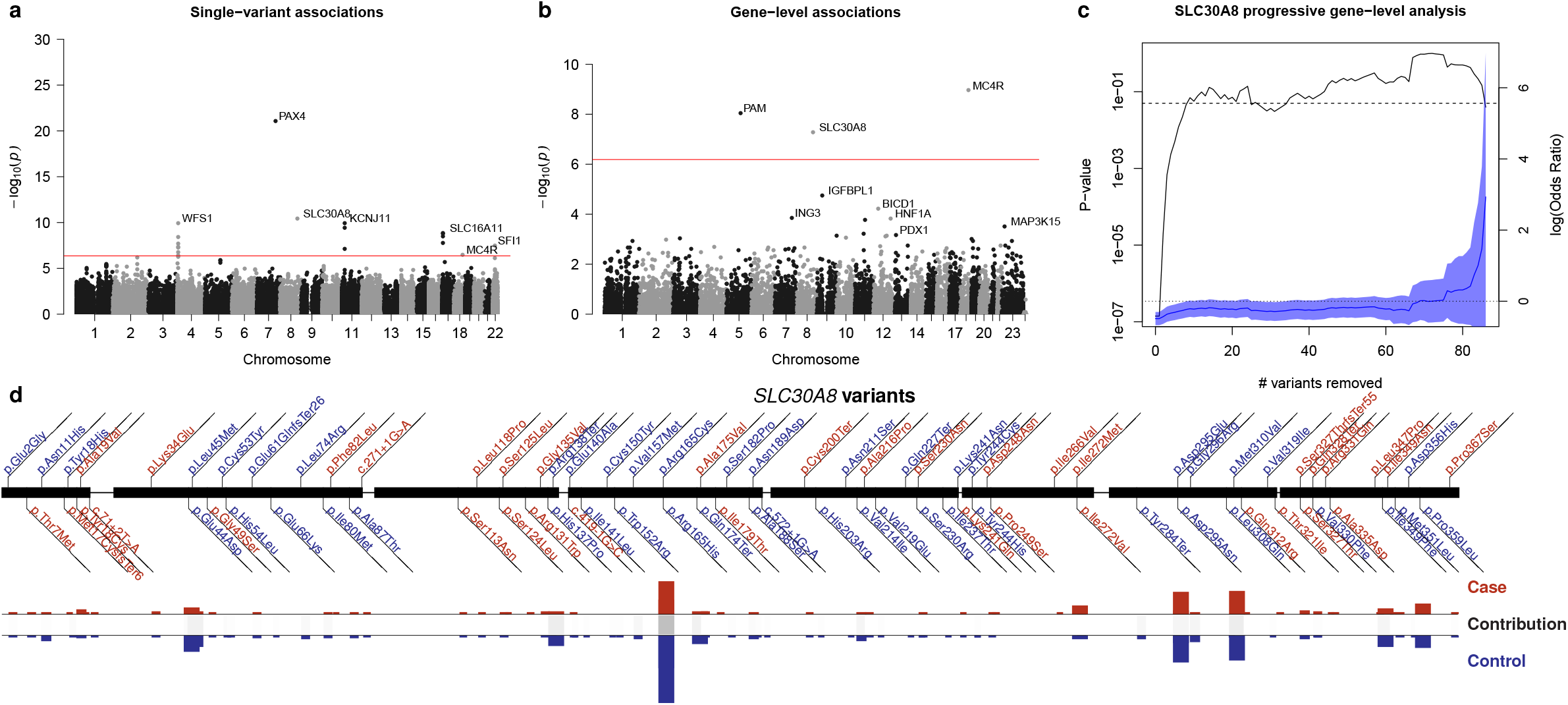
Exome-wide association analysis. **(a)** A Manhattan plot of exome sequence single-variant associations. Genes closest to variants achieving *p*<4.3×10^−7^ (red line; at most one per each 250KB region) are labeled. **(b)** A Manhattan plot of gene-level associations; *p*-values shown are the minimum across the four gene-level analyses after correction for four analyses (**Methods**), with the most significant genes labeled. Red line: *p*=6.5×10^−7^. **(c)** Gene-level association *p*-values for *SLC30A8*, using the burden test on alleles in the 1/5 1% mask (the mask, as defined in **Methods**, achieving greatest statistical significance for *SLC30A8*), after progressive removal of variants in order of increasing single-variant association *p*-value. The left y-axis (black line) shows the progressive gene-level *p*-value, the dashed line *p*=0.05. The right y-axis (blue line) shows the estimated effect size (log_10_(OR)), with shaded blue indicating the 95% confidence interval and dotted line indicating effect size=0. **(d)** Variants observed in *SLC30A8* within 1/5 1% mask. Variants are colored blue (if OR < 1) or red (OR > 1). Case (red) and control (blue) frequencies are shown for each variant, with black boxes shaded according to the contribution of each variant to the gene-level signal (computed by the difference in log_10_(*p*-value) after removal of the variant from the test). OR: odds ratio.

*MC4R* p.Ile269Asn was the sole variant with association OR>2 (Hispanic/Latino MAF=0.89%; *p*=3.4×10^−7^, OR=2.17 [95% CI: 1.63-2.89]). *MC4R* has long established effects on body-weight and diabetes^32–34^, and p.Ile269Asn specifically has been shown to decrease MC4R activity^35,36^ with associations to obesity and T2D in smaller studies of a United Kingdom family^37^ and a Native American population^36^.

As single-variant analysis has limited power to detect associations with rarer variants^12^, we next performed tests of association for sets of variants within genes. We performed two gene-level association tests: (a) a burden test, which assumes all analyzed variants within a gene are of the same effect, and (b) SKAT^38^, which allows variability in variant effect size (and direction).

Following previous studies^22–24^, we separately tested seven different “masks” of variants grouped by similar predicted severity. As this analysis strategy led to 2×7=14 *p*-values for each gene, we developed two methods to consolidate these results for each test (**Methods; Supplementary Figures 8-10**). First, we retained only the smallest *p*-value but corrected for the effective number of independent masks tested^39^, on average 3.6 per gene (“minimum *p*-value test”). Second, we tested all nonsynonymous variants (i.e. missense, splice site, and protein truncating) but weighted each variant according to its estimated probability of causing gene inactivation^12^ (“weighted test”, in essence assessing the effect of gene haploinsufficiency from combined analysis of protein-truncating and missense variants; **Methods**). We verified that the minimum *p*-value and weighted consolidation methods were both well-calibrated (**Supplementary Figure 11**) and between them produced broadly consistent but distinct results: across the ten most significantly-associated genes, *p*-values were nominally significant under both methods for eight genes but varied by one-to-three orders of magnitude (**Supplementary Table 5**). We employed a conservative Bonferroni-corrected gene-level exome-wide significance threshold of *p*=0.05/(2 tests × 2 consolidation methods × 19,020 genes)=6.57×10^−7^.

Using this strategy, gene-level associations reached exome-wide significance for *MC4R, SLC30A8*, and *PAM* (**Figure 1b, Supplementary Tables 5-6**). All three genes lie within previously T2D GWAS loci and contain previously identified coding single-variant signals: p.Arg325Trp and a series of 12 protective protein truncating variants (PTVs) for *SLC30A8*^19,40^, p.Asp563Gly and p.Ser539Trp for *PAM*^24,41^, and p.Ile269Asn for *MC4R*.

In addition to 11 previously observed PTVs, the *SLC30A8* gene-level signal includes 92 variants (103 in total with combined MAF=1.4%; p.Arg325Trp was not included in gene-level analysis) and is associated with T2D protection (weighted *p*=1.3×10^−8^, OR=0.40 [0.28-0.55]). Many variants contributed to this signal: when we progressively removed variants with the smallest single-variant *p*-values, removal of 33 was required to extinguish nominal (*p*<0.05) gene-level significance (**Figure 1cd, Supplementary Figure 12**). Although *SLC30A8* (and its protein product ZnT8) were first implicated in T2D over a decade ago^40^, their molecular disease mechanism(s) remain poorly understood^42,43^ – in part because of seemingly conflicting observations of the common risk-increasing allele p.Arg325Trp (suggested to decrease protein activity^44^) and the rare risk-decreasing PTVs (also thought to decrease protein activity^19^). The protective allelic series from our analysis argues that decreased, rather than increased, risk is the more typical effect of *SLC30A8* genetic variation, and it further provides many alleles that could be characterized to offer mechanistic insight.

The *MC4R* (combined MAF=0.79%; minimum *p*=2.7×10^−10^, OR=2.07 [1.65-2.59]) and *PAM* (combined MAF=4.9%; weighted p=2.2×10^9^, OR=1.44 [1.28-1.62]) gene-level signals are due largely – but not entirely – to effects from individual variants (p.Ile269Asn for *MC4R*, p.Asp563Gly and p.Ser539Trp for *PAM*). For *MC4R*, gene-level association decreased but remained significant after removing p.Ile269Asn (*p*=8.6×10^−3^; **Supplementary Figure 13**). Similarly, as shown previously^34,45^, association was less significant after conditioning on sample BMI, both for the p.Ile269Asn single-variant signal (*p*=1.0×10^−5^) and the gene-level signal not attributable to p.Ile269Asn (*p*=0.035).

The gene-level signal in *PAM* also remained nominally significant (*p*<0.05) even after removing the 35 strongest individually associated *PAM* variants, indicating a contribution from substantially more variants than p.Asp563Gly and p.Ser539Trp (**Supplementary Figure 14**). Cellular characterization of p.Asp563Gly and p.Ser539Trp recently identified a novel mechanism for T2D risk through altered insulin storage and secretion^46^. Our results provide many more genetic variants – identifiable only through sequencing^17^ – that could be characterized for further insights into the T2D risk mechanism mediated by *PAM*.

We finally assessed the 50 most-significant gene-level associations (as measured by minimum *p*-value across our four analyses; **Methods**) in two independent exome sequence datasets: 14,118 individuals (3,062 T2D cases and 9,405 controls of European or African-American ancestry) from the CHARGE discovery sequence project^47^ (CHARGE, **Supplementary Table 7**; 50 genes available) and 49,199 individuals (12,973 T2D cases and 36,226 controls of European ancestry) from the Geisinger Health System (GHS, **Supplementary Table 8**; 44 genes available). In each replication study, *MC4R, SLC30A8*, and *PAM* all showed burden test associations directionally consistent with those from our analysis. *MC4R* (minimum *p*=0.0058) and *SLC30A8* (minimum *p*=0.043) further demonstrated nominally significant associations in the GHS burden analysis, and *MC4R* (minimum *p*=0.026) achieved nominal significance in the CHARGE SKAT analysis. The weaker associations in the replication studies compared to our study (**Supplementary Tables 7** and **8**) could be due to a winner’s curse effect combined with differences in procedures for variant calling, quality control, annotation, and association testing.

More broadly, across the genes with replication results available and with burden *p*<0.05 in our analysis, we observed an excess of directionally consistent burden test associations (31 of 46 in CHARGE, one-sided binomial *p*=0.013; 23 of 40 in GHS, one-sided binomial *p*=0.21; overall one-sided binomial *p*=0.011; **Supplementary Table 9**). Future studies may therefore enable several more of the top gene-level signals from our analysis to reach exome-wide significance.

### Further insights from gene-level analysis

*SLC30A8, MC4R*, and *PAM* illustrate how exome-wide significant gene-level associations provide allelic series that could be characterized for pathogenic insights into previously T2D-associated but still incompletely understood genes. We next investigated the utility of less significant gene-level associations to either (a) genetically prioritize genes with no prior evidence of T2D association, (b) predict the effector gene at established T2D GWAS loci, or (c) predict whether loss or gain of protein function increases disease risk. We conducted this analysis at the level of 16 sets of genes connected to T2D from different evidence sources (e.g. genes harboring diabetes-associated Mendelian or common variants, T2D drug targets^48^, or genes implicated in diabetes-related phenotypes from mouse models^49^; **Supplementary Table 10; Methods**).

First, for each gene set, we asked whether its genes had more significant gene-level associations than expected by chance. We used a one-sided Wilcoxon Rank-Sum Test to compare gene-level *p*-values within each gene set to those for random sets of genes with similar numbers of variants and aggregate frequencies (**Methods**). Twelve of the 16 gene sets achieved *p*<0.05 set-level associations (**Figure 2a-e, Supplementary Figure 15**), including those for T2D drug targets (*p*=6.1×10^−3^) and for genes reported from mouse models of non-autoimmune diabetes (*p*=5.2×10^−3^) or impaired glucose tolerance (*p*=7.2×10^−6^). Following a previous study that retrospectively validated drug targets from the genetic effects of PTVs^27^, these results demonstrate the value of gene-level associations to prioritize candidate genes – e.g. those that emerge from high-throughput experimental screens^50,51^ – for further investigation. Our study emphasizes the added power of including missense variants in this analysis: set-level *p*-values from analysis of PTVs alone were *p*>0.05 for almost all gene sets (although, notably, the drug target gene set remained significant at *p*=0.0061; **Supplementary Figure 16**).

**Figure 2:**
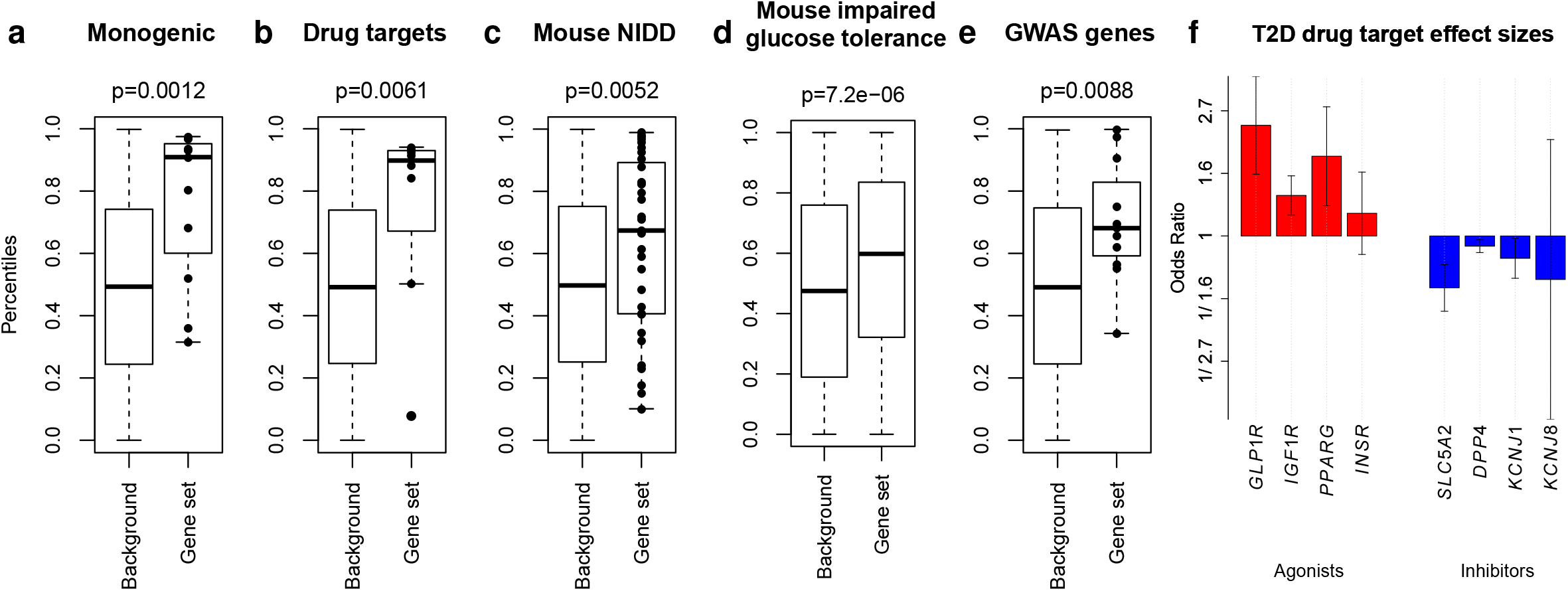
Gene set analysis. **(a-e)** Box plots of the rank percentiles (1 being the highest) for gene-level associations within **(a)** 11 genes implicated in Maturity Onset Diabetes of the Young (MODY); **(b)** 8 genes annotated in the DrugBank database as the primary targets of T2D medications; **(c)** 31 genes annotated in the Mouse Genome Informatics (MGI) database as harboring knockout mutations causing non-insulin dependent diabetes; **(d)** 323 genes annotated in the MGI database as harboring knockout mutations causing impaired glucose tolerance in mice; and **(e)** 11 genes with strong genetic evidence for harboring common causal coding variants. *P*-values correspond to a one-sided Wilcoxon Rank-Sum test comparing the associations to those of matched comparison genes. **(f)** Estimated odds ratios (OR) of deleterious nonsynonymous variants in the eight T2D drug targets. Targets of agonists are colored red and targets of inhibitors are colored blue. Error bars indicate one standard error.

Next, we investigated whether effector genes that mediate GWAS associations – which mostly correspond to variants of uncertain regulatory effects – were also enriched for coding variant gene-level associations. We tested for associations within two sets of predicted effector genes: a curated list of 11 genes harboring likely causal common coding variants (reported from a recent study^17^ with posterior probability of causal association >0.25 from genetics alone; **Methods**), and 20 genes significant in a transcript association analysis with T2D^52^. Genes with likely causal coding variants demonstrated a significant set-level association relative to comparison gene sets (*p*=8.8×10^−3^) and to genes within the same loci (*p*=0.028; **Figure 2e**), even when we conditioned gene-level associations on all significant common variant signals. Most of this signal was due to the gene-level *SLC30A8* and *PAM* associations (*p*=0.082 for the other nine genes). By contrast, the transcript-association based gene set did not exhibit a significant association (*p*=0.72).

Extending this analysis, we curated a list of 94 T2D GWAS loci, and 595 genes that lay within 250 kb of any T2D GWAS index variant, from a 2016 T2D genetics review^53^. Among these 595 genes, 40 achieved a *p*<0.05 gene-level signal (**Supplementary Table 11**), greater than the 595×0.05=29.75 expected by chance (*p*=0.038). These 40 genes had among them significantly more indirect protein-protein interactions (DAPPLE^54^ *p*=0.03; observed mean=11.4, expected mean=4.5) than did the 184 genes implicated based on proximity to GWAS tag SNPs (DAPPLE *p*=0.64), consistent with a gene set of greater biological coherence. Rare coding variants could therefore, in principle, complement common variant fine mapping^6,55^ and experimental data^7,56^ to help interpret T2D GWAS associations, although our results indicate that much larger sample sizes will be required to clearly implicate specific effector genes.

Finally, we assessed whether gene-level analysis could help predict whether gene inactivation increases or decreases T2D risk (i.e. the T2D “directional relationship”^18,19^). For each gene set, we compared the ORs estimated from gene-level weighted analysis of predicted damaging coding alleles (**Methods**) to directional relationships previously reported. Gene-level ORs were 100% concordant with the known relationships for the set of eight T2D drug targets (4/4 inhibitor targets OR<1, 4/4 agonist targets OR>1; one-sided binomial *p*=3.9×10^−3^; **Figure 2f**).

Conversely, concordances between gene-level OR estimates and mouse knockout observations were more equivocal (7/11 diabetes genes with OR>1, binomial *p*=0.27; 137/240 increased circulating glucose genes with OR>1, *p*=0.016; **Supplementary Figure 17**). The relatively low concordances for these gene sets, despite a clear trend toward lower-than-expected gene-level *p*-values within them (**Supplementary Figure 15**), highlight how coding variants might be used to assess seemingly promising preclinical results (particularly given the known limitations of animal models^57,58^). For example, the protective gene-level *ATM* signal we observe (burden test of PTVs OR=0.50, p=0.003) questions previous expectations, based on insulin resistance and impaired glucose tolerance in *Atm* knockout mice^59^, that *ATM* loss-of-function should increase T2D risk. Evidence is even less favorable that *ATM* haploinsufficiency strongly increases T2D risk, rejecting (for example) OR>2 at *p*=1.3×10^−8^. This observation could be relevant in the ongoing characterization of *ATM* as a potential metformin target^60–62^ or if *ATM* activators are considered to treat cardiovascular disease^63^.

### Comparison of rare and common variants in T2D genetic analyses

The substantial number of rare coding variant T2D associations we observed prompted us to re-evaluate arguments^13,14,16,64^ about their value in genetic studies relative to common variants, which have the advantage of being efficiently studied (in many more samples than currently can be sequenced) through array-based association studies^55,65^. While recent studies have emphasized the main contribution of common variants to T2D heritability^17,21,24,66^, they have lacked power to fully evaluate the relative merits of rare versus common variants (or, by implication, sequencing versus array-based studies) to discover disease-associated loci, explain disease heritability, or elucidate allelic series.

For a fair comparison of discoveries possible from sequencing and array-based studies, we collected genome-wide array data within the same individuals we sequenced (available for 34,529 [76.3% of] individuals; 18,233 cases and 17,679 controls). We then imputed variants using best-practice reference panels^67,68^ and conducted single-variant analysis following the same protocol as for the sequence data (“imputed GWAS”; **Supplementary Table 12, Methods**). Eight of the ten exome-wide significant nonsynonymous single-variant associations from our sequence analysis were detectable in the imputed GWAS analysis, together with genome-wide significant noncoding variant associations in 14 additional loci (**Figure 3a, Supplementary Table 13**). All ten single-variant sequence associations were also present on the Illumina Exome Array (**Methods**), implying the ability of array-based association studies to detect exome-wide significant single-variant associations at equivalent significance and at far lest cost than exome sequence association studies.

**Figure 3:**
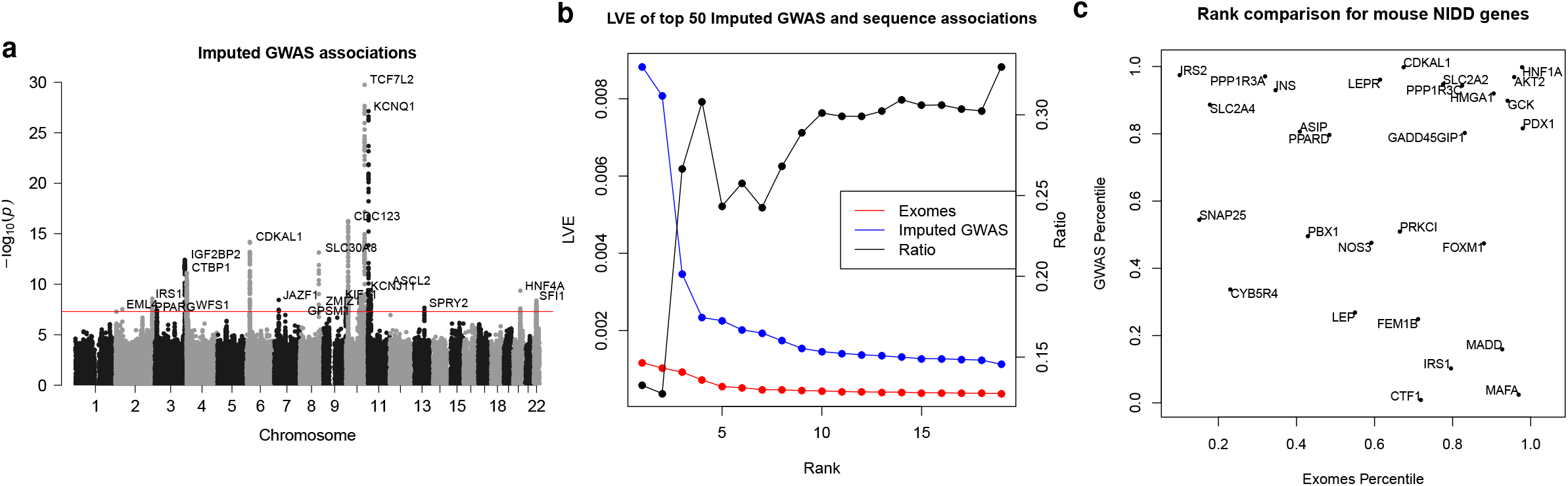
Comparison of exome sequencing to array-based GWAS. **(a)** A Manhattan plot of single-variant associations in an array-based imputed GWAS of the subset (76%) of the samples in the exome sequence analysis for which array data were available. Labels and y-axis are equivalent to **Figure 1a**. **(b)** The observed liability variance explained (LVE) by the top 19 gene-level associations from the exome sequence analysis (red; Exomes) and the top 19 single-variant associations (considering only one per 250kb) from the imputed GWAS (blue; Imputed GWAS), as well as their ratio (black; Ratio). Signals are ranked by LVE rather than *p*-value. **(c)** A comparison of gene rank percentiles according to exome sequence gene-level analysis (x-axis) and gene rank percentiles according to proximity to GWAS signals from a published transethnic T2D GWAS (y-axis; **Methods**). Genes shown are from the set of 31 genes implicated in non-insulin dependent diabetes from knockout mice (the set in **Figure 2c**).

We next compared the contributions to T2D heritability from the strongest (common) single-variant associations from the imputed GWAS to those from the strongest (mostly rare variant) gene-level associations from the sequence analysis. Using a genetic liability model^69^ in which all damaging variants in a gene have the same direction of effect (**Methods**), the three exome-wide significant gene-level signals explain an estimated 0.11% (*MC4R*), 0.092% (*PAM*), and 0.072% (*SLC30A8*) of T2D genetic variance. These estimates are only 10-20% of the variances explained by the three strongest independent common variant associations in the imputed GWAS of the same samples (*TCF7L2*, 0.89%; *KCNQ1*, 0.81%; and *CDC123*, 0.35%) and if anything overstate the heritability explained by rare variants in the gene-level signals, since the *MC4R* and *PAM* estimates are attributable mostly to the low-frequency p.Ile269Asn (70.9% of the gene-level total) and p.Asp563Gly (83.3%) alleles. We obtained similar results in a broader comparison between all (19) previously identified index SNPs achieving *p*<5×10^−8^ in the imputed GWAS and the top 19 gene-level signals from our sequence analysis (**Figure 3b**).

These results argue against a large contribution to T2D heritability from rare variants in the strongest observed gene-level signals, with one caveat: as gene-level tests may include benign alleles that can dilute evidence for association, their aggregate effects might underestimate the true contribution of rare functional variants to T2D heritability^12^. However, when we analyzed all possible subsets of variation in the three most significant gene-level signals (**Methods**), none explained more than 20% of the heritability of the single-variant *TCF7L2* association (maximum of 0.18% for *MC4R*, 0.15% for *PAM*, 0.17% for *SLC30A8*).

We finally assessed whether an array-based study could have detected the allelic series we observed from exome sequence analysis. Among the variants contributing to the exome-wide significant gene-level associations in *SLC30A8, MC4R*, and *PAM*, 95.3% were not imputable (r^2^>0.4; **Methods**) from the 1000 Genomes multiancestry reference panel^67^, and 74.6% of those in Europeans were not imputable from the larger European-focused Haplotype Reference Consortium panel^68^. Similarly, 90.2% of variants (79.7% of European variants) are absent from the Illumina Exome Array.

Additionally, gene set associations using gene “scores”^70^ (**Methods**) from imputed GWAS associations were suggestive (four gene sets achieving *p*<0.05, nine achieving *p*<0.1; **Supplementary Figure 18**) but weaker than gene set associations from our sequence analysis. Some of these gene set associations can be recaptured in larger array-based studies: scores from a published multi-ancestry GWAS of ~110K samples produced *p*<0.05 for 12 of the 16 gene sets we studied (**Supplementary Figure 19, Methods**). However, even here the genes (and corresponding variants) responsible for the gene set associations were broadly different between the array and sequence-based studies, as the two methods often produced uncorrelated rank-orderings of genes within gene sets (e.g. r=−0.11, *p*=0.57 for the mouse diabetes gene set; **Figure 3c**). Collectively, these results argue that array-based GWAS and exome sequencing are complementary, favoring locus discovery and enabling full enumeration of potentially informative alleles, respectively.

### Use of nominally significant associations in translational decision support

The T2D drug targets we analyzed exemplify the opportunities and challenges of using current exome sequence datasets in translational research. Gene-level associations are significant across these targets as a set (**Figure 2b**), and rare variants predict the correct disease directional relationship for each gene (**Figure 2f**). However, rare variant gene-level signals for these genes are nowhere near detectable at exome-wide significance in our current sample size: 80% power would require 110,000-180,000 sequenced cases (220,000-360,000 exomes in a balanced study, equivalent in effective sample size to 750,000-1,200,000 exomes from a population with T2D prevalence 8%; **Figure 4a**).

**Figure 4:**
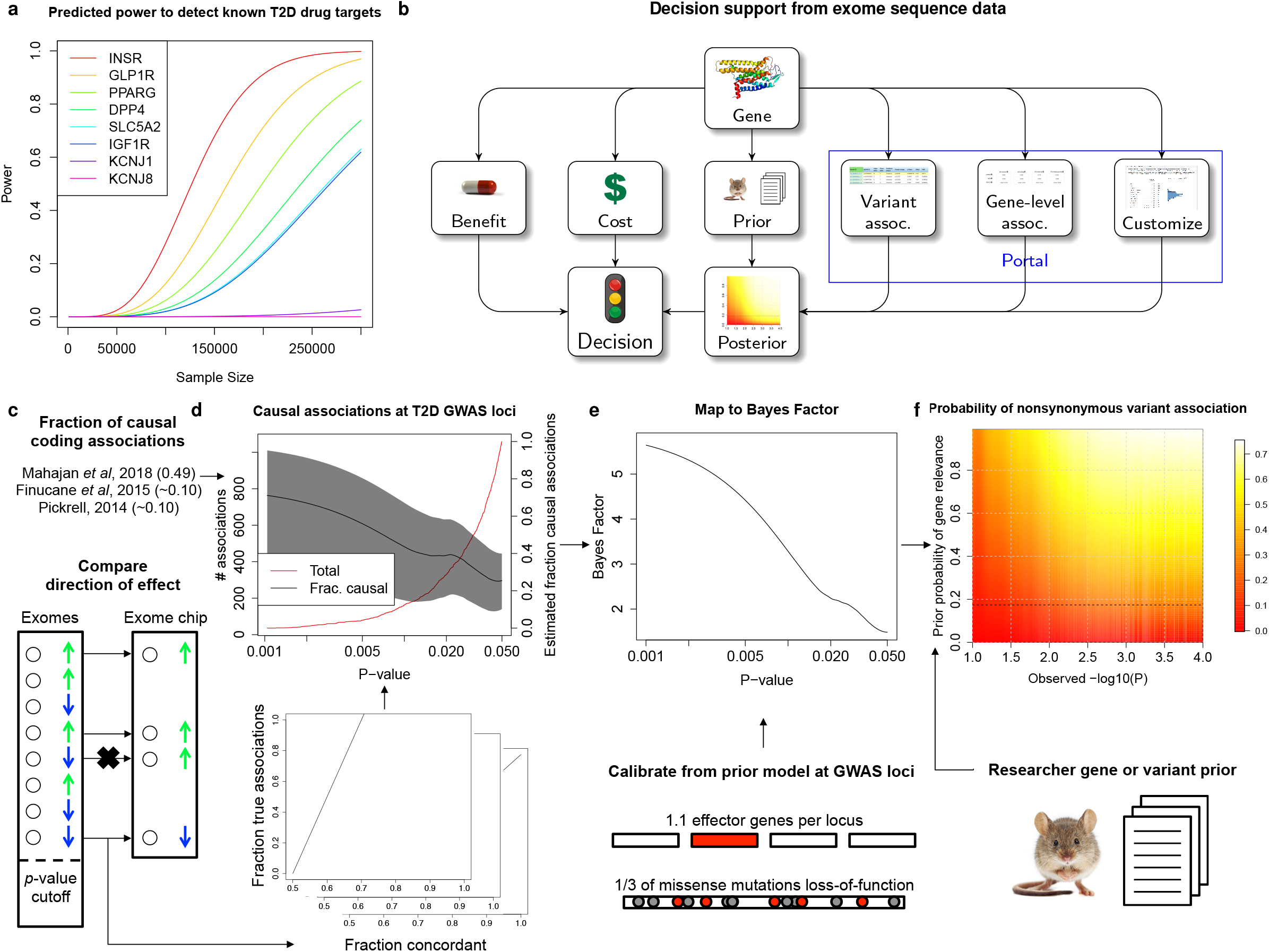
Translational decision support from exome sequence data. **(a)** Estimated power, as a function of future sample size, to detect T2D gene-level associations (at significance *p*=6.25×10^−7^) with aggregate frequency and odds ratios equal to those estimated from our analysis in eight established T2D drug targets (**in Figure 2f**). **(b)** A proposed workflow for using exome sequence data in gene characterization. Depending on the prior belief in the disease-relevance of the gene, the cost of experimental characterization, and the benefit of validating the gene, a decision to conduct a further experiment could be informed by the probability that the gene is relevant to disease, as estimated from exome sequence association statistics (available through www.type2diabetesgenetics.org). **(c-f)** To support this workflow, we estimated the posterior probability of true and causal association (PPA) for nonsynonymous variants in our sequence analysis based on (c) concordance with independent exome chip data and published estimates of the fraction of causal coding associations (**Methods**). **(d)** PPA estimates for nonsynonymous variants within T2D GWAS loci are shown as a function of *p*-value (right y-axis, black; 95% confidence interval, gray) together with the total number of such variants (left y-axis, red). For variants outside of T2D GWAS loci, we developed a method to further compute **(e)** Bayes factors, which measure the odds of true and causal association, as a function of *p*-value, using a model of the prior odds of true and causal association for variants in GWAS loci (**Methods**). These Bayes factors can be **(f)** combined with a subjective prior belief in the T2D-relevance of a gene (y-axis) to produce the estimated posterior probability of true and causal association for any nonsynonymous variant in the exome sequence dataset based on its observed log_10_(*p*-value) (x-axis). Posterior estimates are shaded proportional to value (red: low; white: high). Values shown are for the default modeling assumptions of 33% of missense variants causing gene inactivation and 30% of true missense associations representing the causal variant.

Consequently, many of the more modest associations (e.g. *p*=0.05) in current sample sizes may in fact point to therapeutically relevant variants or genes (**Supplementary Figure 20**)^71,72^. If the false positive rate for these associations – which is expected to be greater than that for associations exceeding exome-wide significance^71–73^ – can be quantified^74,75^, then a modest association signal may motivate further experimentation on a gene while complete absence of an association may reduce enthusiasm for its study. For example, the expected value of the experiment can be calculated based on the likelihood of true association, the cost of the experiment, and the benefit of its success^76,77^ (**Figure 4b**).

We sought to quantify the false positive association rate for nonsynonymous variants observed in our dataset, depending on the *p*-value observed in single-variant analysis. We developed a method to use the consistency of single-variant association statistics between our sequence analysis and a previous^24^ exome array study (re-analyzed to include only the 41,967 individuals not in our current study; **Methods**), together with published estimates of the fraction of nonsynonymous associations that are causal for disease^17,78,79^, to estimate the posterior probability of true and causal association (PPA) for variants reaching different levels of statistical significance. We provide an overview of this method in **Figure 4c-f**, a detailed description in **Methods**, and its sensitivity to modeling assumptions in **Supplementary Figure 21**.

We applied this method to three classes of variants: genome-wide, within T2D GWAS loci, and within genes implicated in T2D through prior (non-genetic) evidence. Model parameters in the middle of the range we explored (**Methods**) predict that 1.5% (95% CI: 0.74%-2.2%) of nonsynonymous variants that achieve *p*<0.05 are truly and causally associated with T2D, increasing to 3.6% (1.4%-5.9%) for variants with *p*<0.005, and 9.7% (3.9%-15.0%) for variants with *p*<5×10^−4^ (**Supplementary Figure 22**). Under this model, 541 (270-810) of the 36,604 nonsynonymous variants with *p*<0.05 in our dataset represent true and causal associations.

Within the set of 94 T2D GWAS loci, we observed evidence of a greater enrichment of true associations: 61.3% of nonsynonymous variants achieving sequence *p*<0.05 were directionally consistent in the independent exome array analysis (compared to 51.9% outside of GWAS loci). We re-calculated a mapping between sequence single-variant *p*-value and PPA using only nonsynonymous variants within these loci. The resulting model predicts that 2.0% (0.048%-4.0%) of such variants overall, 8.1% (3.6%-12.4%) with sequence *p*<0.05, and 17.2% (7.7%-24.1%) with sequence *p*<0.005 represent true and causal T2D associations. This suggests that our dataset contains a large number of potentially strong-effect variants in T2D GWAS loci achieving nominal significance: of 1059 variants with *p*<0.05, we estimate roughly 60 (26-93) of 746 with estimated OR>2 and 41 (18-63) of 503 with estimated OR>3 are true and causal associations (**Supplementary Tables 14-15**).

Beyond GWAS loci, many other genes have evidence – for example from animal^80^ or cellular studies^50,56^ – that may lead a researcher to (often subjectively) believe they are involved in T2D pathogenesis. We extended our approach for PPA estimation to incorporate prior evidence that a gene is relevant to T2D^81^, calibrating it from a model of the prior association likelihood within T2D GWAS loci (**Figure 4e-f; Methods**). Under our model (**Supplementary Table 16**), a prior belief that a gene has (for example) probability 25% of being involved with T2D yields estimates that variants within it achieving *p*<0.05 and *p*<0.005 have 10.7% and 26.2% probabilities of being true and causal T2D associations.

In the future, these PPA calculations could be extended to gene-level associations, which would avoid conflicting results among variants within a gene but require larger-scale gene-level replication data than we had available. Additional work could also develop data and methods to estimate objective, rather than subjective, gene priors and reduce dependence of our conclusions on modeling assumptions (**Supplementary Figure 21**). Still, these PPA calculations provide a useful initial framework to use genetic signals to support cost/benefit estimates of “go/no-go” decisions^82^ in the language of decision theory^76,77^ (**Figure 4b**). To support use of this strategy, we have made our exome sequence association results publically available through the AMP T2D Knowledge Portal (www.type2diabetesgenetics.org), which supports querying of all pre-computed single-variant associations and allows users to dynamically compute single-variant and gene-level associations according to custom covariates and criteria for sample and variant filtering.

## Discussion

Our results paint a nuanced picture of rare variation and T2D, which may also apply to other complex diseases with similar genetic architectures^83^. Our gene set analyses show that rare variant gene-level signals are likely widely distributed across numerous genes, but the vast majority explain, individually, vanishing amounts of T2D heritability – evinced by the >1M samples likely required to detect exome-wide significant rare variant signals in validated therapeutic targets. Gene-level signals that do reach exome-wide significance in our analysis (such as those in *MC4R* and *PAM*) are noteworthy not because they include unusually strong rare variant associations but because they include typical rare variant associations boosted from nominal to exome-wide significance by low frequency variant(s) – which, empirically, can also be detected by array-based studies. Therefore, for many complex traits (particularly those with modest selective pressure like T2D), the primary value of exome sequencing beyond array-based GWAS may be to aid experimental gene characterization^84^ by identifying a broad series of rare coding alleles – ideally through multi-ancestry samples to capture as broad a set of alleles as possible – rather than to discover new disease loci. Whole-genome sequencing will likely, one day, become sufficiently cost effective to subsume both array-based GWAS and exome sequencing; even now, it is at minimum an essential means to expand imputation reference panels to power genetic discovery from GWAS.

Our results also outline a strategy for using exome sequence data to prioritize or validate genes under study by biologists or pharmaceutical industry scientists. We have presented a principled and empirically calibrated Bayesian approach (**Figure 4, Supplementary Table 16**) to estimate the association probability for any variant in our dataset. While currently limited by available data and modeling assumptions, it provides a first step to increase the interpretability of exome sequence associations even absent exome-wide significance. Results and customized analyses from our study can be accessed through a public web portal (www.type2diabetesgenetics.org), advancing the vision to broadly use exome sequence data across many avenues of biomedical research.

## Funding

**Broad Institute, USA**: Sequencing for T2D-GENES cohorts was funded by the National Institute of Diabetes and Digestive and Kidney Diseases (NIDDK) grant U01DK085526: Multiethnic Study of Type Diabetes Genes and National Human Genome Research Institute (NHGRI) grant U54HG003067: Large Scale Sequencing and Analysis of Genomes.

Sequencing for GoT2D cohorts was funded by National Institute of Health (NIH) 1RC2DK088389: Low-Pass Sequencing and High Density SNP Genotyping in Type 2 Diabetes.

Sequencing for ProDiGY cohorts was funded by National Institute of Diabetes and Digestive and Kidney Diseases (NIDDK) U01DK085526.

Sequencing for SIGMA cohorts was funded by the Carlos Slim Foundation: Slim Initiative in Genomic Medicine for the Americas (SIGMA).

Analysis was supported by the National Institute of Diabetes and Digestive and Kidney Diseases (NIDDK) grant U01 DK105554: AMP T2D-GENES Data Coordination Center and Web Portal.

**The Mount Sinai IPM Biobank Program** is supported by The Andrea and Charles Bronfman Philanthropies.

The **Wake Forest study** was supported by NIH R01 DK066358.

**Oxford** cohorts and analysis is funded by: The European Commission (ENGAGE: HEALTH-F4-2007-201413); MRC (G0601261, G0900747-91070); National Institutes of Health (RC2-DK088389, DK085545, R01-DK098032, U01-DK105535); Wellcome Trust (064890, 083948, 085475, 086596, 090367, 090532, 092447, 095101, 095552, 098017, 098381, 100956, 101630, 203141)

The **FUSION** study is supported by NIH grants DK062370 and DK072193.

The research from the **Korean cohort** was supported by a grant of the Korea Health Technology R&D Project through the Korea Health Industry Development Institute (KHIDI), funded by the Ministry of Health & Welfare, Republic of Korea (grant number: HI14C0060, HI15C1595).

**The Malmö Preventive Project and the Scania Diabetes Registry** were supported by a Swedish Research Council grant (Linné) to the Lund University Diabetes Centre.

**The Botnia and The PPP-Botnia studies** (L.G., T.T.) have been financially supported by grants from Folkhälsan Research Foundation, the Sigrid Juselius Foundation, The Academy of Finland (grants no. 263401, 267882, 312063 to LG, 312072 to TT), Nordic Center of Excellence in Disease Genetics, EU (EXGENESIS, EUFP7-MOSAIC FP7-600914), Ollqvist Foundation, Swedish Cultural Foundation in Finland, Finnish Diabetes Research Foundation, Foundation for Life and Health in Finland, Signe and Ane Gyllenberg Foundation, Finnish Medical Society, Paavo Nurmi Foundation, Helsinki University Central Hospital Research Foundation, Perklén Foundation, Närpes Health Care Foundation and Ahokas Foundation. The study has also been supported by the Ministry of Education in Finland, Municipal Heath Care Center and Hospital in Jakobstad and Health Care Centers in Vasa, Närpes and Korsholm. The skilful assistance of the Botnia Study Group is gratefully acknowledged.

**The Jackson Heart Study (JHS)** is supported by contracts HHSN268201300046C, HHSN268201300047C, HHSN268201300048C, HHSN268201300049C, HHSN268201300050C from the National Heart, Lung, and Blood Institute and the National Institute on Minority Health and Health Disparities. Dr. Wilson is supported by U54GM115428 from the National Institute of General Medical Sciences.

**The Diabetic Cohort (DC) and Multi-Ethnic Cohort (MEC)** were supported by individual research grants and clinician scientist award schemes from the National Medical Research Council (NMRC) and the Biomedical Research Council (BMRC) of Singapore.

**The Diabetic Cohort (DC), Multi-Ethnic Cohort (MEC), Singapore Indian Eye Study (SINDI) and Singapore Prospective Study Program (SP2)** were supported by individual research grants and clinician scientist award schemes from the National Medical Research Council (NMRC) and the Biomedical Research Council (BMRC) of Singapore.

**The Longevity study at Albert Einstein College of Medicine, USA** was funded by The American Federation for Aging Research, the Einstein Glenn Center, and National Institute on Aging (PO1AG027734, R01AG046949, 1R01AG042188, P30AG038072).

**The TwinsUK study** was funded by the Wellcome Trust and European Community’s Seventh Framework Programme (FP7/2007-2013). The TwinsUK study also receives support from the National Institute for Health Research (NIHR)-funded BioResource, Clinical Research Facility and Biomedical Research Centre based at Guy’s and St Thomas’ NHS Foundation Trust in partnership with King’s College London.

**Framingham Heart Study** is supported by NIH contract NHLBI N01-HC-25195 and HHSN268201500001I. This research was also supported by NIA AG08122 and AG033193, NIDDK U01 DK085526, U01 DK078616 and K24 DK080140, NHLBI R01 HL105756, and grant supplement R01 HL092577-06S1 for this research. We also acknowledge the dedication of the FHS study participants without whom this research would not be possible.

**The Mexico City Diabetes Study** has been supported by the following grants: RO1HL 24799 from the National Heart, Lung, and Blood Institute; Consejo Nacional de Ciencia y Tecnologi’ a 2092, M9303, F677-M9407, 251M, 2005-C01-14502, and SALUD 2010-2151165; and Consejo Nacional de Ciencia y Tecnologi’ a (CONACyT) [Fondo de Cooperacio’n Internacional en Ciencia y Tecnologi’ a (FONCICYT) C0012-2014-01-247974.

The **KARE cohort** was supported by grants from Korea Centers for Disease Control and Prevention (4845–301, 4851–302, 4851–307), and an intramural grant from the Korea National Institute of Health (2016-NI73001-00).

**The Diabetes in Mexico Study** was supported by Consejo Nacional de Ciencia y Tecnología grant number S008-2014-1-233970 and by Instituto Carlos Slim de la Salud, AC.

**The Atherosclerosis Risk in Communities study** has been funded in whole or in part with Federal funds from the National Heart, Lung, and Blood Institute, National Institutes of Health, Department of Health and Human Services (contract numbers HHSN268201700001I, HHSN268201700002I, HHSN268201700003I, HHSN268201700004I and HHSN268201700005I). The authors thank the staff and participants of the ARIC study for their important contributions. Funding support for “Building on GWAS for NHLBI-diseases: the U.S. CHARGE consortium” was provided by the NIH through the American Recovery and Reinvestment Act of 2009 (ARRA) (5RC2HL102419). CHARGE sequencing was carried out at the Baylor College of Medicine Human Genome Sequencing Center (U54 HG003273 and R01HL086694). Funding for GO ESP was provided by NHLBI grants RC2 HL-103010 (HeartGO) and exome sequencing was performed through NHLBI grants RC2 HL-102925 (BroadGO) and RC2 HL-102926 (SeattleGO).

The infrastructure for the Analysis Commons is supported by R01HL105756 (NHLBI, B.M.P.), U01HL130114 (NHLBI, B.M.P.) and 5RC2HL102419 (NHLBI, E.B.).

**The NHLBI Exome Sequencing Project** (ESP) was supported through the NHLBI Grand Opportunity (GO) program and funded through by grants RC2 HL103010 (HeartGO), RC2 HL102923 (LungGO), and RC2 HL102924 (WHISP) for providing data and DNA samples for analysis. The exome sequencing for the NHLBI ESP was supported by NHLBI grants RC2 HL102925 (BroadGO) and RC2 HL102926 (SeattleGO).

This research was supported by the **Multi-Ethnic Study of Atherosclerosis** (MESA) contracts HHSN268201500003I, N01-HC-95159, N01-HC-95160, N01-HC-95161, N01-HC-95162, N01-HC-95163, N01-HC-95164, N01-HC-95165, N01-HC-95166, N01-HC-95167, N01-HC-95168, N01-HC-95169, UL1-TR-000040, UL1-TR-001079, and UL1-TR-001420. The provision of genotyping data was supported in part by the National Center for Advancing Translational Sciences, TSCI grant UL1TR001881, and the National Institute of Diabetes and Digestive and Kidney Disease Diabetes Research (DRC) grant DK063491.

**The San Antonio Mexican American Family Studies (SAMAFS)** are supported by the following grants/institutes. The San Antonio Family Heart Study (SAFHS) and San Antonio Family Diabetes/Gallbladder Study (SAFDGS) were supported by U01 DK085524, R01 HL0113323, P01 HL045222, R01 DK047482, and R01 DK053889. The Veterans Administration Genetic Epidemiology Study (VAGES) study was supported by a Veterans Administration Epidemiologic grant. The Family Investigation of Nephropathy and Diabetes - San Antonio (FIND-SA) study was supported by NIH grant U01 DK57295. The SAMAFS research team acknowledges late Dr. Hanna E. Abboud’s contributions to the research activities of the SAMAFS.

Samples collection, research and analysis from the **Hong Kong Diabetes Register (HKDR)** at the **Chinese University of Hong Kong (CUHK)** were supported by the Hong Kong Foundation for Research and Development in Diabetes established under the auspices of the Chinese University of Hong Kong, the Hong Kong Government Research Grants Committee Central Allocation Scheme (CUHK 1/04C), a Research Grants Council Earmarked Research Grant (CUHK4724/07M), the Innovation and Technology Fund (ITS/088/08 and ITS/487/09FP), and the Research Grants Committee Theme-based Research Scheme (T12-402/13N).

**The TODAY** contribution to this study was completed with funding from NIDDK and the NIH Office of the Director (OD) through grants U01-DK61212, U01-DK61230, U01-DK61239, U01-DK61242, and U01-DK61254; from the National Center for Research Resources General Clinical Research Centers Program grant numbers M01-RR00036 (Washington University School of Medicine), M01-RR00043-45 (Children’s Hospital Los Angeles), M01-RR00069 (University of Colorado Denver), M01-RR00084 (Children’s Hospital of Pittsburgh), M01-RR01066 (Massachusetts General Hospital), M01-RR00125 (Yale University), and M01-RR14467 (University of Oklahoma Health Sciences Center); and from the NCRR Clinical and Translational Science Awards grant numbers UL1-RR024134 (Children’s Hospital of Philadelphia), UL1-RR024139 (Yale University), UL1-RR024153 (Children’s Hospital of Pittsburgh), UL1-RR024989 (Case Western Reserve University), UL1-RR024992 (Washington University in St Louis), UL1-RR025758 (Massachusetts General Hospital), and UL1-RR025780 (University of Colorado Denver). The content is solely the responsibility of the authors and does not necessarily represent the official views of the National Institutes of Health.

## Acknowledgements

Ruth Loos is supported by the NIH (R01DK110113, U01HG007417, R01DK101855, R01DK107786).

Andrew P Morris is supported by the NIH-NIDDK (U01 DK105535); and a Wellcome Trust Senior Fellow in Basic Biomedical Science (award WT098017).

Jose C Florez is an MGH Research Scholar and is supported by NIDDK K24 DK110550.

Graeme I Bell is supported by P30 DK020595.

Michigan State University is supported by NIH Grant 1K23DK114551-01.

Mark I McCarthy is a Wellcome Trust Senior Investigator (WT098381); and a National Institute of Health Research (NIHR) Senior Investigator. The views expressed in this article are those of the author(s) and not necessarily those of the NHS, the NIHR, or the Department of Health.

Yoon Shin Cho acknowledged support from the National Research Foundation of Korea (NRF) grant (NRF-2017R1A2B4006508).

Ching-Yu Cheng is supported by Clinician Scientist Award (NMRC/CSA-SI/0012/2017) of the Singapore Ministry of Health’s National Medical Research Council.

**LuCAMP**: We wish to thank A. Forman, T. H. Lorentzen and G. J. Klavsen for laboratory assistance, P. Sandbeck for data management, G. Lademann for secretarial support, and T. F. Toldsted for grant management. This project was funded by the Lundbeck Foundation and produced by The Lundbeck Foundation Centre for Applied Medical Genomics in Personalised Disease Prediction, Prevention, and Care (www.lucamp.org). The Novo Nordisk Foundation Center for Basic Metabolic Research is an independent Research Center at the University of Copenhagen partially funded by an unrestricted donation from the Novo Nordisk Foundation (www.metabol.ku.dk). Further funding came from the Danish Council for Independent Research Medical Sciences. The Inter99 was initiated by Torben Jørgensen (principal invesitigator [PI]), Knut Borch-Johnsen (co-PI), Hans Ibsen, and Troels F. Thomsen. The steering committee comprises the former two and Charlotta Pisinger. The study was financially supported by research grants from the Danish Research Council, the Danish Centre for Health Technology Assessment, Novo Nordisk, the Research Foundation of Copenhagen County, the Ministry of Internal Affairs and Health, the Danish Heart Foundation, the Danish Pharmaceutical Association, the Augustinus Foundation, the Ib Henriksen Foundation, the Becket Foundation, and the Danish Diabetes Association. Daniel Witte is supported by the Danish Diabetes Academy, which is funded by the Novo Nordisk Foundation.

We thank all study participants of the Diabetic Cohort (**DC**), Multi-Ethnic Cohort (**MEC**), Singapore Indian Eye Study (**SINDI**) and Singapore Prospective Study Program (**SP2**) for their contributions and the National University Hospital Tissue Repository (**NUHTR**) for biospecimen sample storage.

We thank the Jackson Heart Study (**JHS**) participants and staff for their contributions to this work.

This study was provided with biospecimens and data from the Korean Genome Analysis Project (4845-301), the Korean Genome and Epidemiology Study (4851-302), and the Korea Biobank Project (4851-307, KBP-2013-11 and KBP-2014-68) that were supported by the Korea Centers for Disease Control and Prevention, Republic of Korea.

The **Pakistan Genomic Resource** (PGR) would like to thank all the study participants for their participation. PGR is funded through endowments awarded to CNCD, Pakistan.

The **KORA** study was initiated and financed by the Helmholtz Zentrum München—German Research Center for Environmental Health, which is funded by the German Federal Ministry of Education and Research (BMBF) and by the State of Bavaria. Furthermore, KORA research was supported within the Munich Center of Health Sciences (MC-Health), Ludwig-Maximilians-Universität, as part of LMUinnovativ. For this publication, biosamples from the KORA Biobank as part of the Joint Biobank Munich (JBM) have been used.

Ronald C Ma and Juliana C Chan acknowledged support from the Hong Kong Research Grants Council Theme-based Research Scheme (T12-402/13N), Research Grants Council General Research Fund (Ref. 14110415), the Focused Innovation Scheme, the Vice-Chancellor One-off Discretionary Fund, the Postdoctoral Fellowship Scheme of the Chinese University of Hong Kong, as well as the Chinese University of Hong Kong-Shanghai Jiao Tong University Joint Research Collaboration Fund. We would also like to thank all medical and nursing staff of the Prince of Wales Hospital Diabetes Mellitus Education Centre, Hong Kong.

## Author Contributions

**Leadership**. J.F., N.P.B., J.C.F., M.I.M., M.B. **Analysis team**. J.M.M., C.F., M.S.U., A.Mahajan, T.W.B., L.Chen, S.C., A.E., S.Hanks, A.U.J., K.M., A.N., A.J.P., N.W.R., N.R.R., H.M.S., J.M.T., R.P.W., L.J.S., A.P.M. **Project management/Support roles**. L.Caulkins, R.K., M.C. **Data generation**. Broad Genomics Platform. **T2D-GENES**. A.C., R.A.D., S.G., S.Han, H.M.K., B.-J.K., H.A.K., J.K., J.Liu, K.L.M., M.C.N., M.P., R.S.V., C.S., W.Y.S., C.H.T., F.T., B.T., R.M.v.D., M.V., T.-Y.W., G.Atzmon, N.B., J.B., D.W.B., J.C.C., E.Chan, C.-Y.C., Y.S.C., F.S.C., R.D., B.G., J.S.K., S.H.K., M.L., D.M.L., E.S.T., J.T., J.G.W., E.Bottinger, J.C., J.D., P.F., M.Y.H., Y.J.K., J.-Y.L., J.Lee, R.L., R.C.M., A.D.M., C.N.P., K.S.P., A.R., D.S., X.S., Y.Y.T., C.L.H., G.Abecasis, G.I.B., N.J.C., M.S., R.S., J.B.M., D.A. **GoT2D**. V.L., L.L.B., L.G., P.N., T.D.S., T.T., K.S.S. **LuCAMP**. M.E.J., A.L., D.R.W., N.G., T.H., O.P. **ProDiGY**. L.D., K.L.D., M.K., E.M.-D., C.P., N.S., B.B., P.Z., D.D. **SIGMA**. C.C.-C., E.Córdova, M.E.G.-S., H.G.-O., J.M.M.-H., A.M.-H., E.M.-C., C.R.-M., C.Gonzalez, M.E.G., C.A.A.-S., C.H., B.E.H., L.O., T.T.-L. **CHARGE**. J.W., E.Boerwinkle, J.A.B., J.S.F., N.L.H.-C., C.-T.L., A.K.M., A.C.M., B.M.P., S.W., P.S.d.V., J.D., S.R.H., C.J.O’D., J.P., J.B.M. **Regeneron**. T.M.T., J.B.L., A.Marcketta, C.O’D., D.J.C., H.L.K., F.E.D., A.B., D.C. **KORA**. T.M.S., C.Gieger, T.M., K.S. **ESP**. E.Boerwinkle, M.G., N.L.H.-C., A.C.M., W.S.P., B.M.P., A.P.R., R.P.T., C.J.O’D., L.L., S.R., J.I.R.

## Disclosures

Philip Zeitler is a consultant for Merck, Daichii-Sankyo, Boerhinger-Ingelheim, and Janssen.

Bruce M Psaty serves on the DSMB of a clinical trial funded by Zoll LifeCor and on the Steering Committee of the Yale Open Data Access Project funded by Johnson & Johnson.

## Methods

### Sample selection

We drew samples for exome sequencing from six consortia (**Supplementary Table 1**):

1. The T2D-GENES (Type 2 Diabetes Genetic Exploration by Next-generation sequencing in multi-Ethnic Samples) consortium, an NIDDK-funded international research consortium seeking to identify genetic variants for T2D through multiethnic sequencing studies^24^.
2. The Slim Initiative in Genomic Medicine for the Americas: Type 2 Diabetes (SIGMA T2D), an international research consortium funded by the Carlos Slim Foundation to investigate genetic risk factors of T2D within Mexican and Latin American populations and translate those finding to improved methods of treatment and prevention^85^.
3. The Genetics of Type 2 Diabetes (GoT2D) consortium, an NIDDK-funded international research consortium seeking to understand the allelic architecture of T2D through low-pass whole-genome sequencing, deep exome sequencing, and high-density SNP genotyping and imputation^24^.
4. The Exome Sequencing Project (ESP), an NHLBI-funded research consortium to investigate novel genes and mechanisms contributing to heart, lung, and blood disorders through whole exome sequencing^86^.
5. The Lundbeck Foundation Centre for Applied Medical Genomics in Personalised Disease Prediction, Prevention, and Care (LuCamp) study, which researches whole exome variation in Danish metabolic diseases including diabetes^21^.
6. The ProDiGY (Progress in Diabetes Genetics in Youth) consortium, an NIDDK-funded research consortium to investigate genetic variants for childhood T2D.

Each consortium provided individual-level information on T2D case-control status according to study-specific criteria as well as key covariates including age, sex, and BMI (**Supplementary Table 1**). In addition, several consortia provided data on fasting glucose, 2-hour glucose following glucose challenge, and use of anti-hyperglycemic medications. We excluded as controls individuals with a 2-hour glucose value ≥ 11.1 mmol/L (which meets diagnostic criteria for T2D) or with any two of the following features suggestive of T2D: fasting glucose ≥ 7 mmol/L, hemoglobin A1c ≥ 6.5%, or recorded as taking an anti-hyperglycemic medication. We opted to require two of the previous features since there is room for error in each: fasting values used in T2D diagnostic criteria are required to represent at least an eight-hour fast, accuracy varies across hemoglobin A1c assays, and anti-glycemic medications are occasionally taken by non-diabetic individuals.

All samples were approved for use by their home institution’s institutional review board or ethics committee, as previously reported^21,24,85,86^. Samples newly sequenced at The Broad Institute as part of T2D-GENES, SIGMA, and ProDiGY are covered under Partners Human Research Committee protocol # 2017P000445/PHS “Diabetes Genetics and Related Traits”.

Availability of sequence data and phenotypes for this study is available via the database of Genotypes and Phenotypes (dbGAP) and/or the European Genome-phenome Archive, as indicated in **Supplementary Table 1**.

### Sample Sequencing

For roughly half the study participants (some of T2D-GENES^24^, GoT2D^24^, SIGMA-T2D^85^, LuCAMP^21^, ESP^86^), exome sequence data were available from previous studies. For these individuals (**Supplementary Table 1**), we obtained access to and aggregated BAM files containing unaligned sequence reads, which were generated and analyzed as previously described^23,62,79,80^.

For the remaining participants, de-identified DNA samples were sent to the Broad Institute in Cambridge, MA, USA where samples with (a) sufficient total DNA quantity and minimum DNA concentrations (as estimated by Picogreen) and (b) high quality genotypes (as measured by a 24 SNP Sequenom iPLEX assay) were advanced for subsequent sequencing. Library construction was performed as previously described^87^ with some slight modifications. Initial genomic DNA input into shearing was reduced from 3μg to 50ng in 10μL of solution and enzymatically sheared. For adapter ligation, dual-indexed Illumina paired end adapters were replaced with palindromic forked adapters with unique 8 base index sequences embedded within the adapter and added to each end.

In-solution hybrid selection was performed using the Illumina Rapid Capture Exome enrichment kit with 38Mb target territory (29Mb baited), including 98.3% of the intervals in the Refseq exome database. Dual-indexed libraries were pooled into groups of up to 96 samples prior to hybridization, with liquid handling automated on a Hamilton Starlet Liquid Handling system. The enriched library pools were quantified via PicoGreen after elution from streptavidin beads and then normalized to a range compatible with sequencing template denature protocols.

Following sample preparation, the libraries prepared using forked, indexed adapters were quantified using quantitative PCR (KAPA Biosystems), normalized to 2 nM, and pooled by equal volume using the Hamilton Starlet. Pools were then denatured using 0.1 N NaOH. Denatured samples were diluted into strip tubes using the Hamilton Starlet.

Cluster amplification of the templates was performed according to the manufacturer’s protocol (Illumina) using the Illumina cBot. Flowcells were sequenced on HiSeq 4000 Sequencing-by-Synthesis Kits, then analyzed using RTA2.7.3.

### Variant calling and quality control

Sequencing reads for all samples (both newly sequenced and previously sequenced) were processed and aligned to the human genome (build hg19) using the Picard (broadinstitute.github.io/picard/), BWA^88^, and GATK^89^ software packages, following best-practice pipelines; data from previously published studies were treated the same as data from the new study (i.e. beginning from unaligned reads) to ensure uniformity of processing. Single nucleotide and short indel variants were then called using a series of GATK commands (version nightly-2015-07-31-g3c929b0): ApplyRecalibration, CombineGVCFs, CombineVariants, GenotypeGVCFs, HaplotypeCaller, SelectVariants, and VariantFiltration. Variants were called within 50bp of any region targeted for capture in any sequenced cohort.

We computed hard calls (the GATK-called genotypes but set as missing at a genotype quality [GQ] <20 threshold) and dosages (the expected alternate allele count, defined as *Pr*(RX|data) + 2*Pr*(XX|data), where R is the reference allele and X the alternative allele) for each individual at each variant site. We used hard calls for quality control and dosages in downstream association analyses. We computed dosages on the X chromosome (outside of the pseudo-autosomal region) accounting for sex, treating males as haploid.

To perform data quality control, we first calculated a range of metrics measuring sample sequencing quality (**Supplementary Figure 2**). We then stratified samples by ancestry and sequence capture technology and excluded from further analysis samples that were outliers according to any metric, based on visual inspection by comparison to other samples within the same stratum. A full list of metrics used for exclusion and the number of samples excluded based on each metric is shown in **Supplementary Table 2**.

After exclusion of samples, we calculated an additional set of variant metrics and excluded any variant with overall call rate <0.3, heterozygosity of 1, or heterozygote allele balance of 0 or 1 (i.e. 100% or 0% of reads called non-reference for heterozygous genotypes). We intentionally chose these non-stringent initial variant quality-control thresholds due to the heterogeneity of capture and sequencing technologies used in our study; we performed much more stringent variant quality control during single-variant or gene-level association analysis. We refer to the 49,484 samples and 7.02M variants passing this first round of non-stringent quality control as the “clean” dataset.

### Additional quality control for association analysis in sequence data

Following initial sample and variant quality control, we performed additional exclusions of samples from association analysis. First, we computed a transethnic set of “ancestry” SNPs for use in identity-by-descent (IBD) and principal component (PC) analysis. We began this analysis with variants in the clean dataset (a) with genotype call rate >95%, (b) with minor allele frequency (MAF) >1% in each ancestry, and (c) further than 250Kb from the HLA region or an established T2D association signal. We LD-pruned variants using PLINK^90^ based on maximum r^2^=0.2 (parameters –indep-pairwise 50 5 0.2). We used the remaining 171K variants to estimate pairwise individual IBD using PLINK, and the top 10 PCs of genetic ancestry using EIGENSTRAT^91^. For each pair of individuals with IBD>0.9, we excluded the individual with the lower call rate (337 duplicate exclusions in **Supplementary Figure 2**). We then excluded, for each of the five ancestries, any individual who appeared, based on visual inspection of the first two transethnic PCs, to lie outside of the main PC cluster corresponding to that ancestry (133 ethnic outliers in **Supplementary Figure 2**). Finally, we used the subset of transethnic ancestry SNPs on the X chromosome to compare genetic sex to reported sex, using PLINK, and excluded all discordant individuals (273 sex discordances in **Supplementary Figure 2**).

At this stage we also excluded the 3,510 childhood diabetes cases from the SEARCH and TODAY studies. We initially hoped to include these samples as cases in both single-variant and gene-level analysis, using either PCs or linear mixed models to adjust for any ancestry differences between them and the other samples. However, while single-variant association statistics (computed via a meta-analysis of ancestry-level associations) remained well-calibrated with these studies included (**Supplementary Figure 23ab**), gene-level analysis yielded a dramatically inflated QQ plot (**Supplementary Figure 23cd**). Exclusion of the SEARCH and TODAY study samples, samples failing quality control, and variants that became monomorphic as a result of these sample exclusions, yielded an “analysis” dataset of 45,231 individuals and 6.33M variants.

After these three rounds of sample exclusions, we identified five sets of ancestry-specific “ancestry” SNPs. We used the same procedure as for the transethnic ancestry SNPs (described above), except that we applied the MAF threshold only within the appropriate ancestry. We used these ancestry SNPs to estimate, for each ancestry, pairwise IBD values, genetic relatedness matrices (GRMs), and PCs for use in downstream association analysis.

Additionally, from the IBD values, we generated a list of unrelated individuals within each ancestry by excluding the individual with the lower call rate in any pair of individuals with IBD>0.3 (leading to 2,157 excluded individuals). The resulting “unrelateds analysis” set consisted of 43,090 individuals (19,828 cases and 23,262 controls) and yielded 6.29M non-monomorphic variants. We used this set of individuals and variants for single-variant and gene-level tests (described below) that required an unrelated set of individuals for analysis.

We carried out power calculations^92^ for single-variant or gene-level tests assuming a disease prevalence of 0.08 to convert population frequencies and ORs to case and control frequencies, and a sample size (19,828 cases and 23,262 controls) from an analysis of only unrelated individuals. Our power calculations assumed that allelic effects were homogeneous across ancestries.

### Variant annotation

We annotated variants with the ENSEMBL Variant Effect Predictor^93^ (VEP, version 87). Annotations were produced for all ENSEMBL transcripts with the –flag-pick-allele option used to assign a “best guess” annotation to each variant according to the following ordered criteria for transcripts^94^: transcript support level (TSL, i.e. supported by mRNA), biotype (i.e. protein_coding), APPRIS isoform annotation (i.e. principal), deleteriousness of annotation (i.e. prefer transcripts with higher impact annotations), CCDS^95^ status of transcript (i.e. a high-quality transcript set), canonical status of transcript, and transcript length (i.e. longer preferred). We used the VEP LofTee (https://github.com/konradjk/loftee) and dbNSFP (version 3.2)^96^ plugins to generate additional bioinformatic predictions of variant deleteriousness; from the dbNSFP plugin, we took annotations from 15 different bioinformatic algorithms (listed in **Supplementary Figure 8**) as well as the recent mCAP^97^ algorithm. As these annotations were not transcript-specific, we assigned them to all transcripts for the purpose of downstream analysis.

All single-variant analyses reported in the manuscript or figures are shown using the “best guess” annotation for each variant (as described above).

### Single-variant association analysis in sequence data

To perform single-variant association analysis, we stratified samples by cohort of origin and sequencing technology (i.e. samples from the same cohort but sequenced at different times were analyzed separately). Samples from the ESP study were treated differently, due to the large number of cohorts and sequencing technologies within the study; we stratified ESP samples by ancestry (rather than cohort) and did not further stratify them by sequencing technology. This procedure yielded 25 distinct sample subgroups (**Supplementary Figure 6**).

We then excluded variants separately for each subgroup, based on subgroup-specific measures of call rate, Hardy-Weinberg equilibrium (HWE), differential case-control missingness, and alternate allele genotype quality. Specific filters used to exclude variants from all subgroups are shown in **Supplementary Figure 6**; in general, filters were strict – particularly for multiallelic variants and X-chromosome variants.

For some subgroups, we used stricter filters on top of the basic filters if subgroup-specific quantile-quantile (QQ) plots showed an excess of significant associations. In particular, the Ashkenazi subgroup from the T2D-GENES study showed minimum heterogeneity in sequencing quality between cases and controls (owing to resequencing performed subsequent to the original study publication) and required significant filters to remove artifactual associations. In addition, due to a significant imbalance between the number of cases and controls in the ESP studies, we excluded any variants from that subgroup which had an association *p*-value less than 0.3 times the *p*-value from Fisher’s exact test (under the assumption that covariates in the analysis were inducing statistical artifacts). The numbers of variants passing these filters in each subgroup are shown in **Supplementary Figure 6**.

For each of the 25 sample subgroups, we conducted two single-variant association analyses. In both single-variant analysis, we collapsed all non-reference alleles at multiallelic sites into a single “non-reference” allele.

First, we analyzed all (including related) samples via the EMMAX test^98^, as implemented in the EPACTS (genome.sph.umich.edu/wiki/EPACTS) software package, using the GRM computed from the ancestry-specific ancestry variants. We included in the model covariates for sequencing technology (where appropriate) but not for PCs of genetic ancestry. We did not include covariates for age, sex, or BMI.

Second, we analyzed unrelated samples via the Firth logistic regression test^99^, also as implemented in EPACTS; we included in the model covariates for sequencing technology and for PCs of genetic ancestry (computed from the ancestry-specific ancestry variants). The number of PCs we included varied by subgroup; to select the PCs to be included, we regressed T2D status on sequencing technology and the first ten PCs and included in the model any PC that demonstrated nominal (p<0.05) association with T2D, as well as all higher-order PCs.

For each of the 25×2=50 single-variant analyses, we inspected QQ plots of variant association statistics and increased the stringency of the variant filters if the distribution of association statistics appeared poorly calibrated. The filters shown in **Supplementary Figure 6** represent the final values at which we arrived.

We then conducted a 25-group fixed-effect inverse-variance weighted meta-analysis for each of the Firth and EMMAX tests, using METAL^100^. We used EMMAX results for association *p*-values and Firth results for effect size estimates. For comparison, we conducted two additional meta-analyses with association Z-scores weighted by (a) sample-size and (b) the number of variant carriers. We found that the sample-size weighted meta-analysis had significantly reduced power to detect association for variants with frequencies that varied widely by sample subgroup; for example, 1,425 East-Asian individuals carried p.Arg192His in *PAX4* (N=6,032; *p*=1.2×10^−21^) compared to only 28 carriers across all other ancestries (N=39,199; *p*>0.2), yielding an inverse-variance weighted meta-analysis p=7.6×10^−22^ and a sample-size weighted meta-analysis p=1.0×10^−6^. By contrast, the number-of-carrier weighted meta-analysis yielded similar results as the inverse-variance weighted meta-analysis. We elected to use the inverse-variance weighted method due to its widespread use^100^. We did not conduct random-effects meta-analyses.

### Replication of rs145181683

To assess whether the rs145181683 variant in *SFI1* (*p*=3.2×10^−8^ in the exome sequence analysis) represented a true novel association, we obtained association statistics from the 4,522 Latinos previously analyzed as part of an 8,214 sample Latino GWAS published by the SIGMA-T2D consortium^101^ who did not overlap with the current study. Based on the odds ratio (1.19) estimated in our analysis and the MAF (12.7%) in the replication sample, power was 91% to achieve *p*<0.05 under a one-sided association test. The observed evidence (*p*=0.90, OR=1.00) did not support rs145181683 as a true T2D association.

### Gene-level analysis

We first filtered variants (or, more accurately, alleles, since in contrast to single-variant analysis, we treated multiallelic variants as collections of independent biallelic variants) according to seven different annotation “masks”, ranked in order of increasing deleteriousness. The strongest mask consisted of alleles predicted to cause loss of function by the LofTee algorithm (https://github.com/konradjk/loftee), while weaker masks also included alleles predicted deleterious by progressively fewer bioinformatic algorithms. Each mask included all alleles in higher ranked masks as well as additional alleles specific to the mask. In the two lowest ranked masks (the 1/5 1% and 0/5 1% masks, which included alleles predicted deleterious by one or zero tools, respectively), we filtered alleles specific to each mask according to allele frequency using a cutoff of MAF=1%, with MAF computed as the maximum MAF across the five ancestries. A full list and definitions of masks are shown in **Supplementary Figure 8**; the criteria listed in the figure are for alleles specific to each mask.

To validate that the severity ordering of masks corresponded to an increasing likelihood that an allele in the mask was deleterious, we used previously published data assessing the extent to which all missense variants in the gene *PPARG* impeded adipocyte differentiation (i.e. were annotated as causing *PPARG* loss of function). These data showed a trend whereby alleles in more severe masks had lower predicted functionality (**Supplementary Figure 9**).

For each mask, we grouped alleles by gene according to VEP annotations of impacted transcript; we assigned variants in transcripts of multiple genes to all such genes. For each gene, we created up to three groupings of alleles, corresponding to different transcript sets of the gene. First, the “best” grouping consisted of alleles in the mask according to the “best guess” allele-level annotations. Second, the “all” grouping consisted of alleles in the mask according to any transcript of the gene. Third, the “filter” grouping consisted of alleles in the mask according to protein-coding transcripts of the gene with TSL<3. For many genes, two or more of these allele groupings were identical.

Additionally, we assigned mask-specific allele weights according to their aggregate predicted deleteriousness. To calculate weights, we used a previously published model^12^ in which missense variants are a mixture of fully benign variants and fully loss-of-function variants, with a parameter 0≤*x*≤1 determining the fraction of loss-of-function variants. We assumed all alleles in the LofTee mask were full loss-of-function variants (*x*=1) and that all synonymous alleles were fully benign (*x*=0). We then calculated the (binned) frequency distribution, truncated at MAF<1%, of biallelic LofTee and biallelic synonymous alleles, using these as reference distributions of the frequency of loss-of-function and benign alleles, respectively. For each mask, we then calculated the binned and truncated frequency distribution for alleles specific to the mask (**Supplementary Figure 10**) and estimated a value for *x* (by enumerating and testing a range of possible values between 0 and 1) that maximized the likelihood of the observed frequency distribution. We then used the estimated values of *x* for allele weights, as shown in **Supplementary Figure 8**. Because each mask consisted not only of alleles specific to the mask but also of alleles present in higher ranked masks, alleles within any given mask had a range of weights.

Prior to running gene-level tests, we performed additional quality control on sample genotypes. For each of the 25 sample subgroups (the same subgroups used for single-variant analysis), we identified all variants with low subgroup-specific call rates, high subgroup-specific deviations from HWE, or high subgroup-specific differences between case and control call rates (specific criteria are shown in **Supplementary Figure 8**). For each variant failing any of these criteria, all genotypes for individuals in the subgroup (regardless of allele) were set as “missing”; for multiallelic variants, all subgroup genotypes were set as missing if any allele failed any quality control criterion.

We then conducted a series of tests across the masks. We used a burden test and SKAT^38^, both as implemented in the EPACTS software package. The burden test assumes that the effect sizes of all analyzed variants are the same, while the SKAT test allows effect sizes to vary^102^. We conducted each test across all unrelated individuals pooled together (i.e. in contrast to single-variant analysis, we performed a “mega-analysis” rather than a meta-analysis) and included ten PC covariates (computed from the transethnic ancestry SNPs) as well as indicator covariates for the 25 sample subgroups (the same as defined in single-variant analysis). We did not include covariates for age, sex, or BMI in our analysis, as they had little effect on our results.

We implemented subgroup-specific genotype filters (as defined in the previous quality control step) by modifying the EPACTS software to set specified genotypes to missing during association testing; we achieved allele-specific tests for multiallelic variants (i.e. in which only one allele was present in the mask) in a similar manner by setting non-reference genotypes to missing for samples that carried an allele outside of the mask. We also modified the EPACTS software to accept allele-specific weights by multiplying genotypes (or more accurately, genotype dosages) by the relevant weight prior to conducting the formal burden or SKAT analysis.

### Consolidation of tests across masks

Historically, exome sequencing studies have produced separate gene-level association results for each allelic mask. While straightforward to report, interpreting multiple *p*-values for each gene can be challenging – particularly if the goal is to determine whether a specific gene demonstrates association with a phenotype. To address this challenge, we developed two methods to collapse association results across different allelic masks.

The first method (“weighted test”) collapses associations under a model whereby the phenotypic effects of alleles are directly proportional to their bioinformatically estimated deleteriousness. In the “weighted burden” test, we used the sum of the weights of alleles carried by an individual as a predictor variable in place of the total number of alleles carried. In the “weighted SKAT” test, we multiplied the default weights used in the SKAT EPACTS implementation by the allelic weights we calculated. For these weighted tests we included all alleles in the 0/5 1% mask in the analysis.

Because bioinformatically predicted severity is an imperfect proxy to actual phenotypic severity, we developed a second method, the “minimum *p*-value test”, to collapse associations across masks. We chose the minimum *p*-value test to provide a principled extension of an *ad hoc* but intuitive way to interpret multiple *p*-values for a given gene: take the smallest *p*-value observed across each mask and then correct for the effective number of tests performed for the gene.

To conduct these minimum *p*-value tests, we first ran the burden and SKAT analyses for each of the seven masks separately, following usual exome sequence analysis protocols by using no weights and including all alleles in each mask. For each gene, we then converted the seven *p*-values into a single *p*-value via the formula

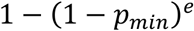

where *e* is the effective number of independent tests performed across the masks. To estimate *e*, we applied a previous approach^39^ originally developed to compute the effective number of independent *p*-values across a set of SNPs:

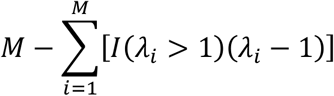

where in our case *M* equals the number of masks (usually seven, except for genes that lack variants in one or more masks or for which two masks are identical) and λ_i_ are the eigenvalues of the *M*×*M* matrix of correlations among the *p*-values of the mask-level tests. To compute the mask *p*-value correlation matrix, we followed the previous approach by first calculating the mask genotype correlation matrix (i.e., for each mask, producing a vector with the number of variants in the mask carried by each individual, and then calculating correlations of the vectors) and then transforming the genotype correlation matrix according to the previously empirically derived^39^ polynomial equation:

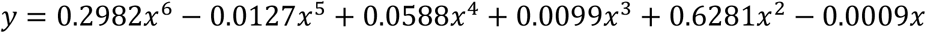

where *x* is the measured correlation between the number of alleles carried and *y* is the estimated correlation between *p*-values.

We note that this polynomial equation was initially developed to translate correlations between individual variants and *p*-values, rather than correlations between aggregate sets of variants and *p*-values, and thus may not be as accurate in our setting. However, genomic control estimates (λ=0.67) and QQ plots (**Supplementary Figure 11**) suggested that if anything our multiple test correction was conservative for most genes. Furthermore, even if our gene-level *p*-values were Bonferroni corrected for all seven masks, the results of our study would remain largely unchanged: each of *SLC30A8, MC4R*, and *PAM* would still exceed exome-wide significance (for both the weighted and minimum *p*-value tests), and the gene set tests would remain nearly identical (as they are based on gene-level *p*-value ranks rather than absolute values). Future work could investigate the application of other methods previously developed to correct for correlated *p*-values^103,104^.

The application of two different methods for collapsing *p*-values across masks for each of two tests yielded four analyses for each gene, corresponding to a weighted burden analysis, a weighted SKAT analysis, an minimum *p*-value burden analysis, and an minimum *p*-value SKAT analysis. In fact, for each of the four analyses, multiple *p*-values were possible for each gene (corresponding to the different transcript sets used for annotation). To produce a single gene-level *p*-value for each of the four analyses, we thus collapsed (for each gene) the set of *p*-values across transcript sets into a single gene-level *p*-value using the same procedure as for the minimum *p*-value test (i.e. taking the minimum *p*-value corrected for the effective number of tests performed).

For some genes (**Supplementary Figures 12-14**) we conducted additional gene-level analyses to dissect the aggregate signals observed. First, we performed tests for each mask separately, including only variants specific to the mask (rather than all variants), to understand whether the aggregate signal was observed in only one as opposed to multiple masks. Second, we performed tests by progressively removing variants in order of lowest single-variant analysis *p*-value, to understand the (minimum) number of variants that contributed statistically to the aggregate signal. Third, we performed tests conditional on each variant separately (i.e. calculating separate models with each individual variant as a covariate), with the resulting *p*-values compared to the full gene-level *p*-value, to assess the contribution of each variant individually to the signal.

### Analysis of exomes from the Geisinger Health System (GHS)

We obtained gene-level association results previously computed from an analysis of 49,199 individuals (12,973 T2D cases and 36,226 controls) from the Geisinger Health System. We requested association summary statistics for the 50 genes with the strongest gene-level associations from our analysis; 44 genes had precomputed summary statistics available; pseudogene *UBE2NL* and X chromosome genes *MAP3K15, SLC16A2, MAGEB5, DGKK*, and *MAGEE2* were not available.

GHS sequence data were processed and analyzed as previously described^27^ and association results were produced for four (nested) variant masks:

1. M1: predicted loss-of-function variants, according to the VEP, with MAF<1% – similar to the LofTee mask but with an additional MAF<1% filter and without the LofTee filter on protein-truncating variants annotated by the VEP.
2. M2: nonsynonymous variants predicted deleterious by 5/5 prediction algorithms with MAF<1% – similar to the 5/5 mask but with an additional filter on MAF<1%.
3. M3: all nonsynonymous variants predicted deleterious by ≥1/5 bioinformatic algorithms with MAF<1% – similar to the 1/5 1% mask.
4. M4: all nonsynonymous variants with MAF<1% – similar to the 0/5 1% mask, although not identical as the 1% filter was used for all variants including those in the LofTee and 5/5 masks.

For each mask, association results were computed via logistic regression under an additive burden model (with phenotype regressed on the number of variants carried by each individual) with age, age^2^, and sex as covariates. Although this analysis procedure was broadly consistent with the one we used for our exome sequence analysis, we were not able to synchronize our procedures for quality control, annotation, and collapsing association statistics across masks.

To produce a single GHS p-value for each gene, we applied the minimum *p*-value procedure across the four mask-level results. We estimated the correlation matrix using the same procedure as for our exome sequence analysis, using the combined GHS allele frequencies reported across the four (nested) masks.

### Analysis of exomes from the CHARGE consortium

We collaborated with the CHARGE consortium to analyze the 50 genes with the strongest gene-level associations from our analysis in 12,467 individuals (3,062 T2D cases and 9,405 controls) from their previously described study^105^. CHARGE DNA samples were processed at Baylor College of Medicine Human Genome Sequencing Center using the VCRome 2.1 design and sequenced in paired-end mode in a single lane on the Illumina HiSeq 2000 or the HiSeq 2500 platform with a mean 78-fold coverage. All samples were called together and details on sequencing, variant calling, and variant quality control were described in detail by Yu et al.^106^

Variants in the CHARGE exomes were annotated and grouped into seven masks using the same procedure as for the original exome sequence analysis. For each mask, CHARGE burden and SKAT association tests were performed in the Analysis Commons^107^ using a logistic mixed model^108^ assuming an additive genetic model and adjusted for age, sex, study, race, and kinship.

To produce a single CHARGE *p*-value for each gene, we applied the minimum *p*-value procedure across the four mask-level results, as for the GHS analysis.

### Evaluation of directional consistency between exome sequence, CHARGE, and GHS analyses

We examined the concordance of direction of effect size estimates (i.e. OR>1 or OR<1) between our original exome sequence analysis and those from CHARGE and GHS. We used burden test statistics for this analysis, as SKAT tests do not produce direction of effects. Of the 50 genes advanced for replication, we considered the 46 that reached burden *p*<0.05 for at least one mask (i.e. ignoring those with evidence for association only under the SKAT model). We compared the direction of effect to that estimated by burden analysis of the same (or analogous) mask in the GHS or CHARGE analysis. For CHARGE, we compared direction of effect for the same mask. For GHS, we compared use the following approximate mapping between masks: LofTee to M1; 15/15, 10/10, 5/5, and 5/5+LofTee LC to M2; 1/5 1% to M3; and 0/5 1% to M4. We then conducted a one-sided exact binomial test to assess whether the fraction of results with consistent direction of effects was significantly greater than expected by chance.

### Generation of candidate T2D-relevant genes sets

To assess whether gene-level association strength could be an informative metric to use when prioritizing candidate genes for further study or experimentation, we compared gene-level associations for genes in a variety of gene sets (**Supplementary Table 10**) to gene-level association statistics for random sets of genes matched with the target set based on the number and frequencies of variants (as described below). We did so for 16 sets of genes:

1. *Eleven genes harboring mutations that cause Maturity Onset Diabetes of the Young (MODY)*. We selected genes from a set previously described^24^ after excluding two genes (*ABCC8* and *KCNJ11*) that can cause monogenic diabetes or congenital hyperinsulinism depending on whether the mutations they harbor are activating or inactivating.
2. *Eight genes annotated as targets for antidiabetic medications*. We downloaded medications annotated as “Drugs Used in Diabetes” or “Blood Glucose Lowering” from the DrugBank database version 5.0^48^. After exclusion of medications with more than two annotated targets, we advanced for analysis only genes (a) annotated as a target of at least two compounds and (b) for which the therapeutic target modulation strategy was consistently annotated across all medications, where annotations of “inhibitor”, “antagonist”, and “inverse agonist” were interpreted as reducing activity, while annotations of “agonist”, “activator”, or “inducer” were interpreted as increasing activity. These restrictions excluded *ABCC8* from analysis, as it was annotated as the target of both an inhibitor and an agonist; we elected to maintain this exclusion, despite multiple lines of evidence^109^ indicating inhibition of ABCC8 to be the appropriate anti-diabetic strategy, to maintain consistent criteria across all genes selected for analysis. Additionally, we excluded *KCNJ11* (which with *ABCC8* encodes the ATP-sensitive K(ATP) channel targeted by sulfonylureas) from analysis because both medications listed in DrugBank as targeting it had more than two targets (Glyburide, 8, and Glimepiride, 3). The resulting gene set was thus *GLP1R, IGF1R, PPARG, INSR, SLC5A2, DPP4, KCNJ1*, and *KCNJ8*.
*3-14. Twelve sets of genes reported as relevant to T2D in mouse models*. Within the Mouse Genome Informatics Database, we searched for genes matching various diabetes-relevant “phenotypes, alleles, and disease models” under the broader category of “mouse phenotypes and mouse models of human disease”. We constructed a gene set for each phenotype defined in the database, many of which overlapped. For phenotypes associated with increased diabetes risk, we used: (3) “type 2 diabetes or type ii diabetes” (i.e. non-insulin dependent diabetes; 31 genes), (4) “diabetes mellitus” (72 genes), (5) “impaired glucose tolerance” (327 genes), (6) “increased circulating glucose” (365 genes), (7) “insulin resistance” (181 genes), and (8) “decreased insulin secretion” (133 genes). For phenotypes associated with decreased diabetes risk, we used: (9) “improved glucose tolerance” (239 genes), (10) “decreased circulating glucose” (481 genes), (11) “increased insulin sensitivity” (178 genes), and (12) “increased insulin secretion” (51 genes). For phenotypes associated with diabetes risk but with unclear direction of effect, we used (13) “decreased circulating insulin” (321 genes) and (14) “increased circulating insulin” (215 genes).
*15. Eleven genes suspected of harboring common coding causal variants within T2D GWAS loci*. We analyzed the set of genes from a recent exome array analysis^17^ which contained a coding variant GWAS signal for which the unweighted posterior probability of causality exceeded 25%. Although the final values reported by the study include an elevated prior for coding variants, we elected to use a 25% unweighted posterior threshold to enrich for the genes with the highest likelihood of mediating the observed GWAS signal. For analysis of this gene set, we recomputed gene-level association statistics within the set by conditioning on all GWAS tag SNPs (within the locus) reported in the exome array analysis^17^; we used *p*-values from these conditional gene-level associations in the gene set analysis.
*16. Twenty genes with T2D-associated transcript levels*. We selected genes with significant associations in a pre-publication^52^ tissue-wide T2D association analysis (i.e. testing for association between the genetic component of tissue-level gene expression and T2D), with associations considered significant if they survived Bonferroni correction for all tested genes and all tested tissues. Results were computed with the MetaXcan software package^110^ using SNP regression coefficients taken from a large trans-ethnic T2D GWAS meta-analysis^111^ and gene expression prediction models from the PredictDB website (http://predictdb.org).

### Gene set analysis

For each gene set, our goal was to compare the gene level *p*-values within the set to those of genes chosen at random from the genome. To control for gene variability in the number and frequency of variants within them, which could confound comparisons, we constructed comparison genes by matching on four properties: the (1) number of variants in any of the seven variant masks; (2) total allele counts over all variants in any of the seven masks; (3) number of tests across all variant masks and transcript sets; and (4) effective number of tests across all variant masks and transcript sets (as computed for the minimum *p*-value test). We scaled each property to zero mean and unit variance. For each gene, we then used the 50 nearest neighbors (defined using Euclidean distance in the scaled property space) as matched comparison genes.

To conduct a gene set analysis, we then combined the genes in the gene set with all of the comparison genes matched to each gene in the set. Within the combined list of genes, we ranked genes using the *p*-values observed for the minimum *p*-value burden test. We then used a one-side Wilcoxon rank-sum test to assess whether genes in the gene set had significantly higher ranks than the comparison genes. For gene set analysis, we used the minimum *p*-value test, rather than the weighted test, under the rationale that (a) we aimed to detect associations with as many genes as possible using information from as many variants as possible and (b) the weighted test might not detect genes that did not follow its model of a strong correlation between variant effect sizes and molecular annotation. We used the burden test rather than SKAT based on a desire to have more interpretable association statistics (e.g. effect size estimates). However, we did not quantitatively and systematically compare the power of each of our analyses in this setting.

### Use of gene-level associations to predict effector genes

In most situations, GWAS associations implicate common regulatory variants, which seldom localize to specific genes. To assess whether gene-level associations from exome sequencing – which are composed mostly of rare variants independent from any GWAS associations – could prioritize potential effector genes within known T2D GWAS loci, we catalogued all genes within each locus reaching *p*<0.05 for the minimum *p*-value burden test. We took a list of 94 GWAS loci from a recent review article^53^ and advanced for analysis the 595 genes within 250kb of an index SNP.

We then sought to compare two methods to predict effector genes within these loci. First, we used *p*<0.05 according to the minimum *p*-value gene-level test from our exome sequence analysis to predict candidate effector genes, producing a list of 40 genes (across 32 loci). Second, we used proximity to the index SNP (as predicted by DAPPLE^54^) to predict candidate effector genes, producing a list of 184 genes (at some loci DAPPLE annotated more than one candidate effector gene).

As accurately assessing which of these two gene sets is more enriched for true effector genes would require (at minimum) significant experimental work, we used the relative number of protein interactions within each gene set as one (imperfect) measure of their respective biological “coherence”. To assess whether each set encodes proteins with more interactions than would be expected by chance, we ran DAPPLE through the public GenePattern portal (https://software.broadinstitute.org/cancer/software/genepattern) with default values for all parameters. The 40 genes with minimum *p*<0.05 were significantly more enriched for protein interactions (*p*=0.03; observed mean=11.4, expected mean=4.5) than were the 184 genes implicated based on proximity to the index SNP (*p*=0.64; observed mean=21.1, expected mean=21.9).

While these results suggest that gene-level associations may be useful for prioritizing effector genes, we note that they do not implicate any specific genes and that DAPPLE is only one means to assess biological coherence of a gene set (through direct and indirect protein interactions). Evaluation of the biological candidacy of these genes may ultimately require in-depth functional studies^56^.

### Use of gene-level associations to predict direction of effect

In therapeutic development, it is often valuable to know the direction of effect linking gene modulation to disease risk – that is, whether inactivation or activation of a protein increases disease risk. We thus assessed whether gene-level association analysis of predicted deleterious variants could be used to predict this direction of effect. For this analysis, we used odds ratios estimated from a modified weighted burden test procedure, which only included alleles from the four masks with the predicted most deleterious variants: LofTee, 16/16, 11/11, and 5/5 (**Supplementary Figure 8**). Weights for variants were identical to those used in the exome-wide weighted burden test. We chose these four masks for analysis to balance a desire for greater aggregate allele count per gene (i.e. missense variants in addition to protein-truncating variants) with a need to strongly enrich for deleterious variants (>73% estimated to be deleterious in masks analyzed vs. <50% in the other masks (**Supplementary Figure 8**). In addition, we used the weighted test because it was explicitly designed to estimate an effect of gene haploinsufficiency based on both protein-truncating and missense variants.

To compare these direction of effect estimates to those expected for T2D drug targets, we assumed agonist targets to have true OR>1 and inhibitors to have true OR<1. For a comparison to expectations for mouse gene knockouts, we first excluded 473 genes annotated, based on membership in multiple gene sets, to have both expected OR>1 and expected OR<1 (these genes were excluded only from the direction of effect comparisons; they were maintained in all other gene set analyses). This left 389 genes with an expected OR>1, associated exclusively with mouse traits indicative of increased risk (overlapping sets of 11 “type 2 diabetes or type ii diabetes”, 46 “diabetes mellitus”, 204 “impaired glucose tolerance”, 245 “increased circulating glucose”, 104 “insulin resistance”, and 63 “decreased insulin secretion”), and 467 genes with an expected OR<1, associated exclusively with traits indicative of decreased risk (overlapping sets of 164 “improved glucose tolerance” genes, 358 “decreased circulating glucose” genes, 95 “increased insulin sensitivity” genes, and 18 “increased insulin secretion” genes). Gene sets for “decreased circulating insulin” and “increased circulating insulin” were excluded from this direction of effect comparison due to the unclear relationship between these phenotypes and T2D risk.

### Aggregation and generation of SNP array data

Because the most significant single-variant associations that emerged from our exome sequence analysis were with common variants, we asked whether an array-based genome-wide association study in the same samples could have provided a less expensive method to detect these same associations. To address this question, we aggregated all available SNP array data for the exome-sequenced samples (**Supplementary Table 12**). Data for the GoT2D^24^, SIGMA^85^, and T2D-GENES consortia have been previously analyzed (unpublished T2D-GENES data were collected from a range of SNP arrays including Affymetrix 5.0 and 6.0, Illumina HumanHap 610K and 1M, and the Illumina CardioMetabochip). The newly sequenced samples from the T2D-GENES and SIGMA consortia were genotyped on a custom “Genomes For Life” (G4L) Illumina Infinium array, including 243,662 variants chosen to uniquely identify each individual in a study and to provide a backbone for imputation of common variation. The G4L array was processed by the Arrays lab of Broad Genomics and called using the Illumina GenCall (Autocall) algorithm.

### Analysis of SNP array data

After genotyping, the 34,529 samples (18,233 cases and 17,679 controls; **Supplementary Table 12**) both in the exome sequence analysis and with a SNP array call-rate >95% were advanced for imputation. To omit variants that might degrade imputation quality, prior to imputation we excluded variants with low genotype call rate (<95%), strong deviation from Hardy-Weinberg equilibrium (*p*<10^−6^), differential genotype call rate between cases and controls (*p*<10^−5^), or low frequency (MAF<1%). We then imputed autosomal variants (SNVs, short indels, and large deletions) via the Michigan Imputation Server^112^ for each of two reference panels: the all ancestries 1000 Genomes Phase 3 (1000G) reference panel of 2,504 individuals^67^ and the Haplotype Reference Consortium (HRC) Panel of 32,470 individuals^68^. We used the 1000G-based imputation for all association analyses and the HRC-based imputation to assess the number of exome sequence variants imputable from the largest available European reference panel. We note that the HRC panel includes only SNPs (i.e. no indels) and only variants observed at least five times in the sequence data contributed to the HRC.

After imputation, we performed sample and variant quality control, as well as association tests, analogous to the exome sequence single-variant analysis. By contrast with the exome sequence analysis, we found that the EMMAX test produced more suspicious looking associations than did the Firth test and thus used only the Firth test (i.e. for both *p*-values and ORs) in the imputed GWAS analysis.

To determine which variants in the exomes dataset were imputable from the 1000G or HRC panel, we calculated which of the exome variants passed imputed GWAS quality control in any sample subgroup, with a further restriction of achieving r^2^>0.4 in that subgroup. Only variants in the exomes dataset that were polymorphic in the imputed GWAS samples were included in this analysis. For calculations involving the HRC-imputed GWAS (given that the HRC panel is European-specific), we only considered variants variable in four European cohorts (METSIM, Ashkenazi, GoDARTS, and FHS) in the analysis.

### Gene set analysis using SNP array data

In addition to single-variant analysis, we conducted gene set analysis with the imputed GWAS data. We first used the method implemented in MAGENTA^70^ to assign gene scores from the imputed GWAS single-variant association results; MAGENTA gene scores are based on proximity to a GWAS lead SNP after correction for potential confounding factors. In the same way as for gene set analysis from the exome sequence gene-level results, we then conducted a one-sided Wilcoxon rank-sum test to compare the gene scores to those of matched comparison genes.

As the imputed GWAS gene set analysis produced fewer significant gene set associations than did the exome sequence gene set analysis, we investigated whether a larger array-based association study would produce more significant gene set associations (i.e. whether the lack of gene set associations in the imputed GWAS was due to a fundamental lack of associated common variants near the genes in the gene set or simply due to an insufficient sample size). For this analysis, we downloaded single-variant association statistics from the largest available multiethnic array-based GWAS for T2D^111^, converted them to MAGENTA gene scores, and then for each gene set conducted a Wilcoxon rank-sum test as described above.

### LVE calculations

To calculate liability variance explained (LVE), we used a previously presented formula^69^ to calculate the LVE of a variant with three genotypes (*AA, Aa*, and *aa*) and corresponding relative risks (1, *RR_1_*, and *RR_2_*). For these calculations we assumed HWE, implying the frequencies of the three genotypes to be *P_aa_*=*P_a_*^2^, *P_Aa_*=2*P_a_*(1-*P_a_*), and *P_AA_*=(1-*P_a_*)^2^, where *P_a_* is the minor allele frequency. Under this assumption, LVE can be expressed as

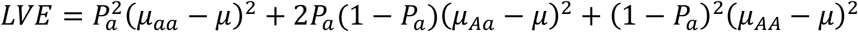

where *μ* = 2*P_a_* (1 − *P_a_*)*μ_Aa_* + (1 − *P_a_*)^2^ *μ_AA_*, and

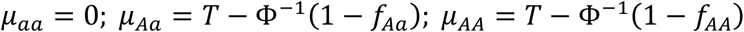

Here Φ^−1^ is the normal quantile distribution, *T* = Φ^−1^(1 − *f_aa_*), and *f_aa_, f_Aa_*, and*f_AA_* are defined as

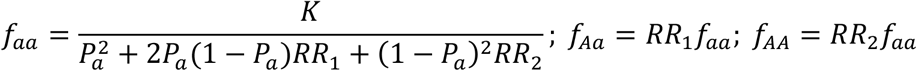

where *K* is the disease prevalence.

The inputs to these formulae are estimates of allele frequency (for either individual variants or sets of variants, depending on whether variant-level or gene-level variance is to be calculated), relative risk, and disease prevalence. For individual variants, we used the point estimate of the MAF from our analysis to estimate allele frequency, while for genes we used the point estimate of combined allele frequency (across all alleles) in place of MAF. We estimated relative risks from analysis ORs and MAFs 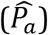 under an assumed prevalence of *K*=0.08 and an additive genetic model, by iteratively solving two equations^69^:

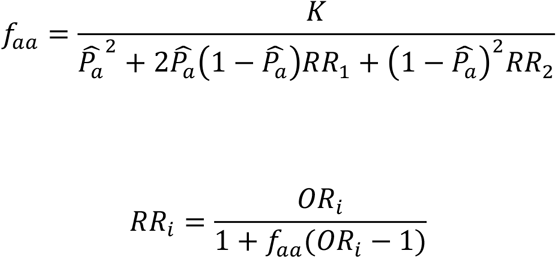

where *i*=1,2 correspond to the heterozygous and major-allele homozygous genotypes. We used a multiplicative model for odds-ratios; i.e. *OR_2_* = *OR_1_^2^*.

We performed LVE calculations as an integral over the distribution of potential relative risks, assuming that the logarithm of odds ratios *OR_i_* followed normal distributions with means and variance equal to those estimated from our analysis. When presenting the strongest LVE values for the imputed GWAS analysis, we only considered variants genotyped in at least 10,000 individuals to avoid potential artifacts resulting from a spurious association in a small sample subgroup.

For gene-level LVE calculations, we used the variant mask with lowest *p*-value to calculate LVE. As each mask may have included a mixture of disease-associated and benign alleles, the calculated LVE may underestimate the true LVE for disease-associated alleles within the gene. To calculate an upper bound on the LVE by only disease-associated alleles, we performed a series of LVE calculations for progressively larger sets of alleles, at each step including alleles by order of decreasing single-variant significance. We performed two calculations for each gene, one for risk alleles and one for protective alleles, taking the maximum of the two as the final upper bound estimated for LVE by the gene. We did not calculate an LVE bound under a model whereby alleles within the gene can both increase and decrease risk of disease.

### Estimated power to detect gene-level associations with T2D drug targets

To estimate the power of future studies to detect gene-level associations in genes with effect sizes similar to those for established T2D drug targets, we used aggregate allele frequencies and odds ratios estimated from our gene-level analysis and an assumed prevalence of *K*=0.08 to calculate a proxy for true population frequencies and relative risks. In each case, we used odds ratios and frequencies from the variant mask yielding the strongest gene-level association. Because on average these drug targets had 5 effective tests per mask, we used an exome-wide significance threshold of *α*=1.25×10^−7^ for power calculations. We calculated power as previously described^92^.

### Estimated fraction of true associations

We sought to quantify the proportion of true associations (PPA) for nonsynonymous variants observed in our dataset as a function of association strength as measured by single-variant *p*-value. We define a true association as a variant which, when studied in larger sample sizes, will eventually achieve statistical significance owing to a true OR≠1. We distinguish *true* association from *causal* association: causally associated variants are the subset of truly associated variants in which the variant itself is causal for the increase in disease risk, as opposed to being truly associated due to LD with a different causally associated variant.

To estimate PPA, we used as training data a previous exome array study from the GoT2D consortium spanning 13 European cohorts^24^. As two of the 13 cohorts included in the previous study contributed samples to the current exome sequence analysis, we re-calculated a fixed-effects inverse-variance weighted meta-analysis for every variant in the exome array study after excluding all samples from these two overlapping cohorts. This yielded a collection of exome array association statistics for 206,373 variants, with a maximum sample size of 50,567 (maximum effective sample size 41,967).

We then compared variant direction of effect estimated from our exome sequence analysis of 45,231 individuals to those estimated from the independent exome array analysis of 41,967 individuals. To produce an uncorrelated set of associations tests for this analysis, we pruned all collections of variants using the LD-clump procedure (parameters –clump-p1 0.1 –clump-p2 0.1 –clump-r2 0.01) of the PLINK software package^90^, which required variants to have pairwise r^2^<0.01. We performed this procedure for (a) nonsynonymous variants within 94 previously established T2D GWAS loci and (b) nonsynonymous variants exome-wide. For the 1,059 nonsynonymous variants within established T2D GWAS loci achieving *p*<0.05 in the exome sequence analysis, the directions of effect were concordant (both OR>1 or both OR<1) with the exome array analysis for 61.3% of variants. This fraction decreased (as expected) for higher *p*-value thresholds (e.g. 49.4% at *p*>0.5) and when only variants outside of T2D GWAS loci were analyzed (51.9% at *p*<0.05).

To estimate the fraction of true associations among the set of variants achieving significance below a threshold *p* (e.g. *p*<0.05), we modeled the set of variants as a mixture of proportions *x_p_* of truly associated variants (OR≠1) and (1-*x_p_*) of truly non-associated variants (OR=1). We assumed non-associated variants have a 50% chance of a concordant direction of effect between the two analyses, and truly associated variants have a greater chance according to their estimated effect size. Specifically, assuming that the observed effect size for a variant follows a normal distribution with mean equal to the true effect and variance that scales inversely with sample size, we estimated the probability *p_i_* of producing a concordant effect for variant v_*i*_ as

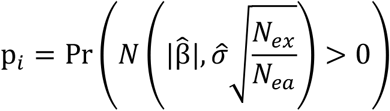

where 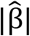 is the absolute value of the estimated (from the exome sequence analysis) logarithm of the odds ratio, 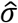 is the estimated standard error of the logarithm of the odds ratio, *N_ex_* is the effective sample size of the exome sequence analysis, and *N_ea_* is the effective sample size of the exome array analysis.

The expected fraction of variants exhibiting concordant direction of effect is then

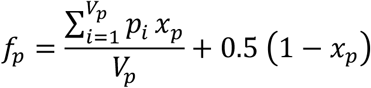

where *V_p_* is the number of variants in the set. Based on the observed fraction 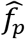 of variants with concordant directions of effect, we thus estimated *x_p_* by

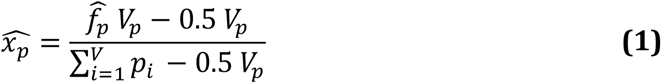

To calculate a 95% confidence interval (CI) for *x_p_*, we first estimated a 95% CI for *f_p_* using the Jeffreys interval method^113^, as implemented in the R software package (https://www.r-project.org), and we then used equation **(1)** to convert its lower and upper bounds to lower and upper bounds on the corresponding confidence interval for *x_p_*.

### Probability of causal association

The estimated values for *x_p_* can be interpreted as estimates of the posterior probability that a variant with *p*<0.05 in our analysis is truly associated with T2D rather than due to chance. As our ultimate goal was to quantify the probability of *causal* association, rather than just true association, we modeled the probability of variant association as a function of (a) the probability of causal association (*PPA_c_*), influenced in turn by the likelihood that the variant results in gene loss-of-function as well as the likelihood that the gene is relevant to T2D; and (b) the prior probability of indirect association (*PPA_i_*), influenced in turn by the likelihood that the variant is in LD with a nearby but different variant that is causally associated with T2D. Under the assumption that causal and indirect associations are disjoint events, this model expresses PPA as

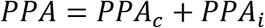

Precisely determining which coding variant associations are in fact causal requires fine mapping of all nearby variants in large sample sizes^6^, which is currently infeasible for the mostly rare variants observed in our study. Since we could not accurately calculate specific values of *PPA_c_* and *PPA_i_* for each variant, we instead used estimates of the average the proportion of associations that are causal (α), where *α* is the probability of causal association *conditional* on a true association, rather than the absolute probability of causal association. We considered two means to estimate α.

First, recent analyses have attempted to assess the contribution of nonsynonymous variants to T2D or similar traits, either by directly estimating the proportion of associations that are due to nonsynonymous variants^79^ or by measuring the proportion of heritability explained by nonsynonymous variants^78^. These analyses suggest that ~10% of T2D associations are likely to be due to nonsynonymous variants. As these calculations apply to all associations in the genome, rather than those in which at least one nonsynonymous variant achieves significance, they likely underestimate the proportion of nonsynonymous associations that are causal.

Second, a recent exome array study identified 40 exome-wide significant nonsynonymous variant associations and then calculated the probability of causal association for each (via credible set analysis)^17^. The reported average probability of causal association across these variants of 49.2% provides a direct estimate of α. This estimate is likely less biased than that based on genome-wide analyses of all T2D associations, but it is based on a small number of associations and thus has a high variance.

Based on these considerations, we considered values of 10%, 30%, and 50% for α. and used 30% as our default value for analyses reported in the main manuscript. For any value of *x_p_*, representing the fraction of true associations at a given *p*-value threshold, we calculated a value for 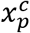, representing the fraction of causal associations at a given *p*-value threshold, as 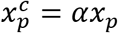. Under this model, using a different value for α (e.g. 50% or 10%) would scale *PPA_c_* estimates linearly (e.g. 5/3 or 1/3 as high).

### Incorporation of prior likelihood into posterior probability estimations

Following previous work^81^, the posterior probability of causal association 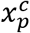 can be expressed as a combination of the prior odds of causal association for the variant, *π* (i.e. the belief, prior to observing any genetic association data, that the variant is causally associated with T2D), and the Bayes factor for causal association of the variant calculated from genetic association data, *BF_c_*:

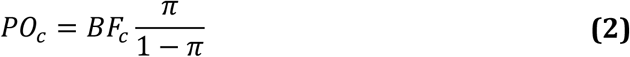

where *PO_c_* is the posterior odds of causal association expressed as

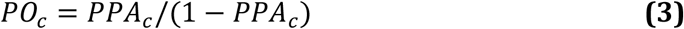

We use a “*c*” subscript in *PO_c_* and *BF_c_* to emphasize that they are posterior odds (and Bayes factors) for causal association, rather than just true association.

Given an estimate 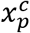 of the posterior probability of causal association (i.e. *PPA_c_*) for a class of variants (e.g. those satisfying *p*<0.05), as well as a prior probability of causal association *π* for the same class of variants, we can calculate an estimate of the average Bayes factor for variants in the class as:

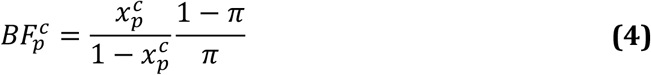

Here, 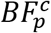 denotes the average Bayes factor for causal association (i.e. the ratio of the likelihood of the observed data under the model of causal association to the likelihood of the observed data under the model of no association) for variants with *p*-value below a given *p*. We note that this equation indirectly infers an average Bayes factor from a direct estimate of an average posterior 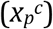 and a specified prior *π*, which is different from how Bayes factors are usually calculated.

Under the assumption that the relationship between a variant’s *π* and *PO_c_* is, given its observed *p*-value, conditionally independent of all other variant properties (i.e. dependence on properties such as sample size is entirely captured by the observed *p*-value), we calibrated the relationship between *p*-value and 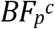 using nonsynonymous variants within GWAS loci. We modeled *π* for such variants assuming (a) on average 1.1 genes within 250kb of each GWAS signal harbors coding variants associated with T2D; (b) missense variants are a mixture of fully benign and fully protein-inactivating variants^12^; (c) only inactivating missense variants; and (d) one-third of missense variants are inactivating (as estimated by the average weight of missense variants in our masks). Based on the 595 genes within the 94 T2D GWAS loci in our analysis, this yielded a prior estimate of 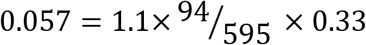.

The gene prior was inspired by the often implicit expectation that a GWAS signal usually represents a single causal variant^114^ affecting a single gene (although multiple effector genes may be more common than previously thought^3^). To assess the sensitivity of our results to the assumption of 1.1 disease-relevant genes per T2D GWAS locus, we repeated all calculations with the additional choices of 0.5 and 2 genes per GWAS locus (**Supplementary Figure 21ab**).

We calculated the variant prior based on the mean weight of variants in our dataset as computed for the “weighted” gene-level test, as these weights were designed to directly estimate the probability that variants in a mask cause full loss of function. This calculation produced a prior estimate of 34.2% for nonsynonymous variants in our dataset, not far from a previously reported value of 25%^12^. We thus used a value of 33% for the variant prior in our main analysis, with values of 40% and 25% used for comparison (**Supplementary Figure 21cd**).

Through the prior probability of causal association for nonsynonymous variants in T2D GWAS loci of 0.057, and equations **(1)**-**(4)** above, we produced a lookup table mapping variant *p*-values to Bayes factors of causal association (*BF_c_*). For any subsequent variant *v* with observed *p*-value *p(v)* and a user-specified prior on the relevance of its gene to T2D, we then calculated its posterior likelihood of association by mapping *p(v)* to *BF_c_* and then employing equations **(2)** and **(3)** to calculate an estimated posterior probability of causal association (*PPA_c_*). Although not presented here, lower and upper confidence intervals on *PPA_c_* can also be estimated by repeating this procedure using the lower and upper confidence intervals for *x_p_^c^* in equation **(4)**.

### Sensitivity of *PPA_c_* to modeling parameters

The above calculations rely on two parameters, the specific values of which will affect final *PPA_c_* estimates. First, they require a parameter for the proportion of true nonsynonymous associations that are causal. As described above and in the text, we used a value – of 30% – in between a published estimate of the proportion of nonsynonymous associations within GWAS loci that are causal (49.2%) and a published estimate of the proportion of causal associations that are nonsynonymous (~10%). Using a different value (e.g. 50% or 10%) would scale the PPAc estimates linearly (e.g. 5/3 or 1/3 as high).

In addition, calculations involving a user-specified prior require a parameter for the proportion of nonsynonymous variants in GWAS loci that causally influence T2D risk (prior to any observed associations). This parameter does not affect *PPA_c_* estimates genome-wide or within GWAS loci, as we directly estimate *PPA_c_* estimates for these genes from our data and therefore do not require a user-specified prior. Although we decompose this parameter into two – a parameter for the proportion of genes within T2D GWAS loci that are relevant to disease and a parameter for the proportion of missense variants within a gene that result in loss of function – only the product of the two parameters is used in the model. **Supplementary Figure 21** shows the impact of different values for these two parameters.

